# CRAG: *De novo* characterization of cell-free DNA fragmentation hotspots in plasma whole-genome sequencing

**DOI:** 10.1101/2020.07.16.201350

**Authors:** Xionghui Zhou, Haizi Zheng, Hailu Fu, Kelsey L. Dillehay McKillip, Susan M. Pinney, Yaping Liu

## Abstract

Non-random cell-free DNA fragmentation is a promising signature for cancer diagnosis. However, its aberration at the fine-scale in early-stage cancers is poorly understood. Here, we developed an approach to *de novo* characterize the cell-free DNA fragmentation hotspots from whole-genome sequencing. In healthy, hotspots are enriched in gene-regulatory elements, including open chromatin regions, promoters, hematopoietic-specific enhancers, and, interestingly, 3’end of transposons. Hotspots identified in early-stage hepatocellular carcinoma patients showed overall hypo-fragmentation patterns compared to healthy controls. These cancer-specific hypo-fragmented hotspots are associated with genes enriched in gene ontologies and KEGG pathways that are related to the initiations of hepatocellular carcinoma and cancer stem cells. Further, we identified the fragmentation hotspots at 297 cancer samples across 8 different cancer types (92% in stage I to III), 103 benign samples, and 247 healthy samples. The fine-scale fragmentation level at most variable hotspots showed cancer-specific fragmentation patterns across multiple cancer types and non-cancer controls. With the fine-scale fragmentation signals alone in a machine learning model, we achieved 48% to 95% sensitivity at 100% specificity in different early-stage cancer. We further validated the model at independent datasets we generated at a small number of early-stage cancers and healthy plasma samples with matched age, gender, and lifestyle. In cancer-positive cases, we further localized cancer to a small number of anatomic sites with a median of 80% accuracy. The results highlight the significance of *de novo* characterizing the cell-free DNA fragmentation hotspots for detecting early-stage cancers and dissection of gene-regulatory aberrations in cancers.

## Introduction

Circulating cell-free DNA (cfDNA) from patients’ plasma is a promising non-invasive biomarker for disease diagnosis[1]. The fragmentation patterns of cfDNA are not evenly distributed in the genome and altered in cancer, bringing enormous signals from both tumor and peripheral immune cells to detect early-stage cancers [2–4]. Recently, several patterns have been derived to capture the full spectrums of the cfDNA fragmentation in cancer, such as patterns near transcription start sites (TSS) and transcription factor binding sites (TFBS), orientation-aware cfDNA fragmentation (OCF), the preferred-ended position of cfDNA, motif diversity score (MDS), large-scale fragmentation patterns at mega-base level (DELFI), nucleosome positioning (window protection score, WPS), and multi-modality integrations[2,5–13]. However, the studies of fragmentation patterns at selected known regulatory elements, such as TSS[6], TFBS[9], and known open chromatin regions in selected immune cells(OCF)[8], limited their opportunities to unbiasedly characterize the genome-wide fragmentation aberrations on other regulatory regions in early-stage cancers. The preferred-ended position of cfDNA has not been associated with known gene-regulatory elements yet[7]. The end motif and MDS[10] is a summary statistic score for each patient that does not allow further explorations of its association with specific gene-regulatory elements. The large-scale fragmentation patterns at mega-bases level (DELFI)[2] are challenging to be associated with the fine-scale gene-regulatory elements, genes, pathways, and therefore further druggable targets for the interventions of early-stage cancers. These challenges limited their potential opportunity to characterize the underlying unknown gene-regulatory aberrations during the initiations of early-stage cancers.

To conquer these challenges, we need an unbiased genome-wide approach to narrow down the regions of interest from cfDNA fragments directly. A previous study on cfDNA from healthy and late-stage cancers *de novo* characterized the regions with high WPS signals that are associated with nucleosome occupancies[5]. Nucleosome occupancies inside the cells are usually measured by MNase-seq, which is not comprehensively performed at various primary cell types across different human pathological conditions, such as cancer. Thus, the characterization of nucleosome-occupied regions from cfDNA, such as WPS, will still limit our scope to dissect the potential regulatory aberrations in cancer. However, the reduced fragmentation process (“fragmentation coldspots”) at nucleosome-occupied regions, on the other side, indicates the potential existence of an increased fragmentation process (“fragmentation hotspots”) in the open chromatin regions. The open chromatin region is a hallmark of DNA regulatory elements and has recently been comprehensively profiled by ATAC-seq and DNase-seq at many primary cell types across different pathological and physiological conditions, including cancer and immune cells[14,15]. Transcription factors, which are critical for disease progression, usually bind the open chromatin regions rather than the nucleosome-occupied regions[16]. Therefore, instead of identifying “fragmentation coldspots” at nucleosome-occupied regions, we hypothesize that the characterization of cfDNA “fragmentation hotspots”, potentially enriched in open chromatin regions and gene-regulatory elements, will not only boost the power for the identification of nuanced pathological conditions, such as early-stage cancer, but also elucidate the unknown *in vivo* gene-regulatory mechanisms indicated by the cfDNA fragmentation patterns from patients’ plasma.

Here, we developed a computational approach, named **C**ell f**R**ee dn**A** fra**G**mentation (CRAG), to *de novo* identify the genome-wide cfDNA fragmentation hotspots by utilizing the weighted fragment coverages from cfDNA paired-end WGS data. We observed the high enrichment of these fragmentation hotspots at open chromatin regions and related gene-regulatory elements. We demonstrated the cfDNA fragmentation aberrations in early-stage cancers. Finally, as a proof-of-concept study, we showed the possibility to utilize these cancer-specific fragmentation hotspots for the detection and localization of multiple cancers, mostly early-stage.

## Results

### CRAG: a probabilistic model to characterize the cell-free DNA fragmentation hotspots

We proposed a computational approach to *de novo* characterize the fine-scale genomic regions with higher fragmentation rates than the local and global backgrounds, defined as cfDNA fragmentation hotspots (Fig. 1a-b). Since both fragment coverages and sizes are essential parts of evaluating the fragmentation process, we weighed the fragment coverages in each region by the ratio of average fragment sizes in the region *versus* that in the whole chromosome, named integrated fragmentation score (IFS) (Details in Methods). The negative binomial model we proposed correctly captured the variation of IFS in the background and indicated the existence of cfDNA fragmentation hotspots (Fig. 1c, Details in Methods). We utilized both local (50kb) and global (whole chromosome) backgrounds to identify the significant hotspots, which is especially useful when the focal copy number changes exist. Since sequencing coverages are usually affected by the G+C% content, we also normalized the IFS signals with the G+C% content within the regions (Details in Methods). We used the cfDNA deep WGS data (BH01, ∼100X)[5] from the healthy non-pregnant individuals as the primary dataset to evaluate our approach in healthy individuals. In the BH01 dataset, we identified 138,938 cfDNA fragmentation hotspots. The IFS distributions in both BH01 and another independent dataset from a healthy individual (IH01, ∼100X) showed expected depletions at the center of BH01 hotspots (Fig. 1d, Fig. S1a), suggesting that we correctly capture the genome-wide fragmentation hotspots.

**Fig. 1.**
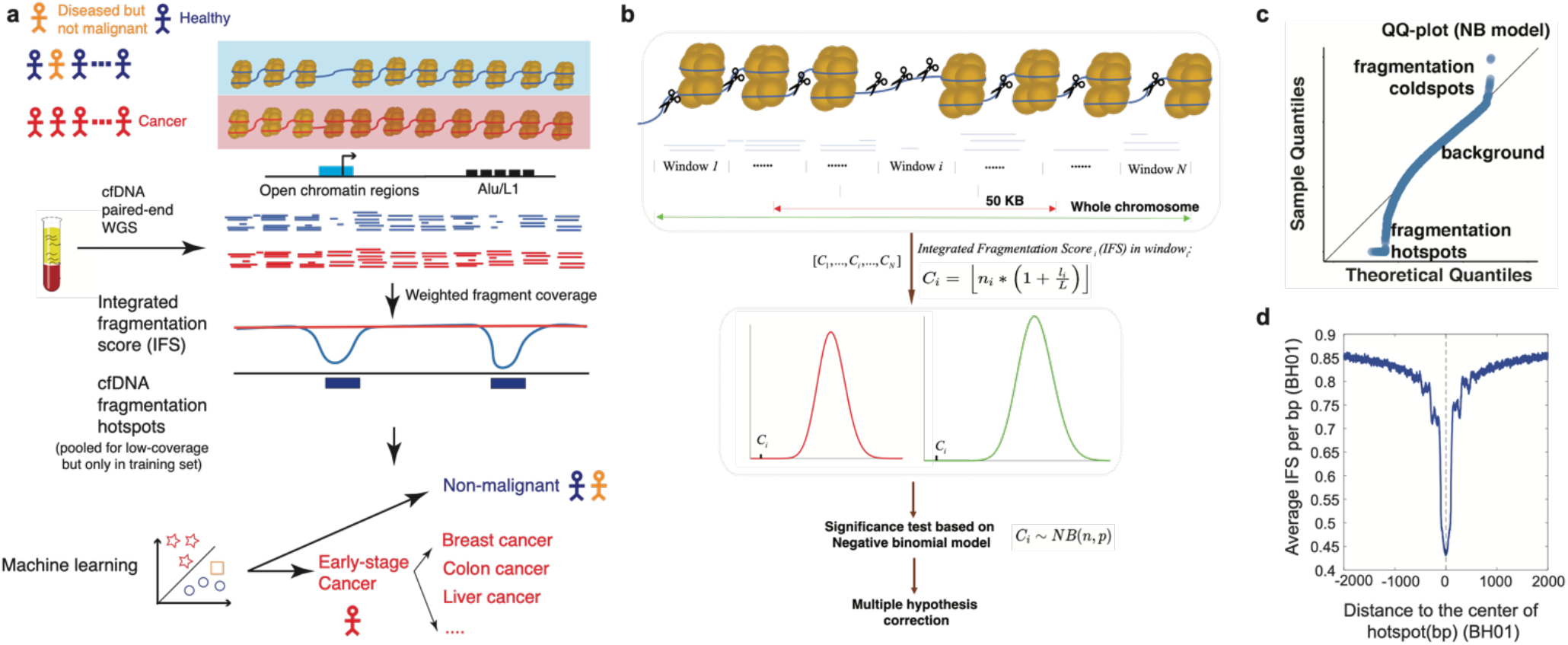
The schematic of CRAG approach. (a). The schematic of using cfDNA hotspots for the cancer diagnosis. (b). The schematic of cfDNA hotspot identification. (c). The Q-Q plot for the negative binomial modeling of IFS score distribution. (d). The distribution of IFS around the hotspots (BH01, healthy).

It is well known that the fragment coverages and lengths from next-generation sequencing, including cfDNA WGS, are affected by the sequence compositions[5,17,18]. To check if the depletion of IFS signals is just due to the bias of sequence compositions, we normalized the IFS signals by k-mer composition (n=2) at BH01 hotspots (Details in Methods). We did not observe any change in the overall distribution of fragmentation patterns before and after the correction (Fig. S1b). These results suggested that our model robustly captured the fragmentation hotspots in cfDNA WGS.

### Cell-free DNA fragmentation hotspots are highly enriched in open chromatin regions and active gene-regulatory elements

We next sought to characterize the genomic distributions of these fragmentation hotspots in healthy individuals (BH01). Similar to the previous studies on the open chromatin regions[19], the fragmentation hotspots from cfDNA are highly enriched at the CpG island (CGI) promoters and CTCF insulators, but not enriched at the non-CGI promoters, 5’ exon boundaries, transcription termination sites (TTS), and random genomic regions (Fig. 2a). We plotted the distributions of publicly available DNA accessibility and active/repressive histone modification marks from the major hematopoietic cell types around the hotspots. We found the high enrichment of epigenetic marks related to the active regulatory element as expected (Fig. 2b-c, Fig. S2-S3). Moreover, the enhancer mark H3K4me1, from hematopoietic cell types but not other cell types, showed a high enrichment around the hotspots, which is consistent with previous studies that hematopoietic cell types are the major contributors to cfDNA in healthy individuals[20–22] (Fig. 2d, Fig. S3). To further understand the enrichment of fragmentation hotspots at different chromatin states, we utilized the 15-states chromHMM segmentation results across different cell types from the NIH Roadmap Epigenomics Mapping Consortium[23]. The hotspots mainly showed the enrichment in the tissue/cell-type-specific chromHMM states from hematopoietic cell types but not other cell types. (Fig. 2e). The evolutionary conservation score (phastCons) in hotspots is significantly higher than in matched random regions (two-sided Mann–Whitney U test, p < 2.2e^-16^, Fig. S4) [24], further suggesting the enrichment of the functional regulatory elements.

**Fig. 2.**
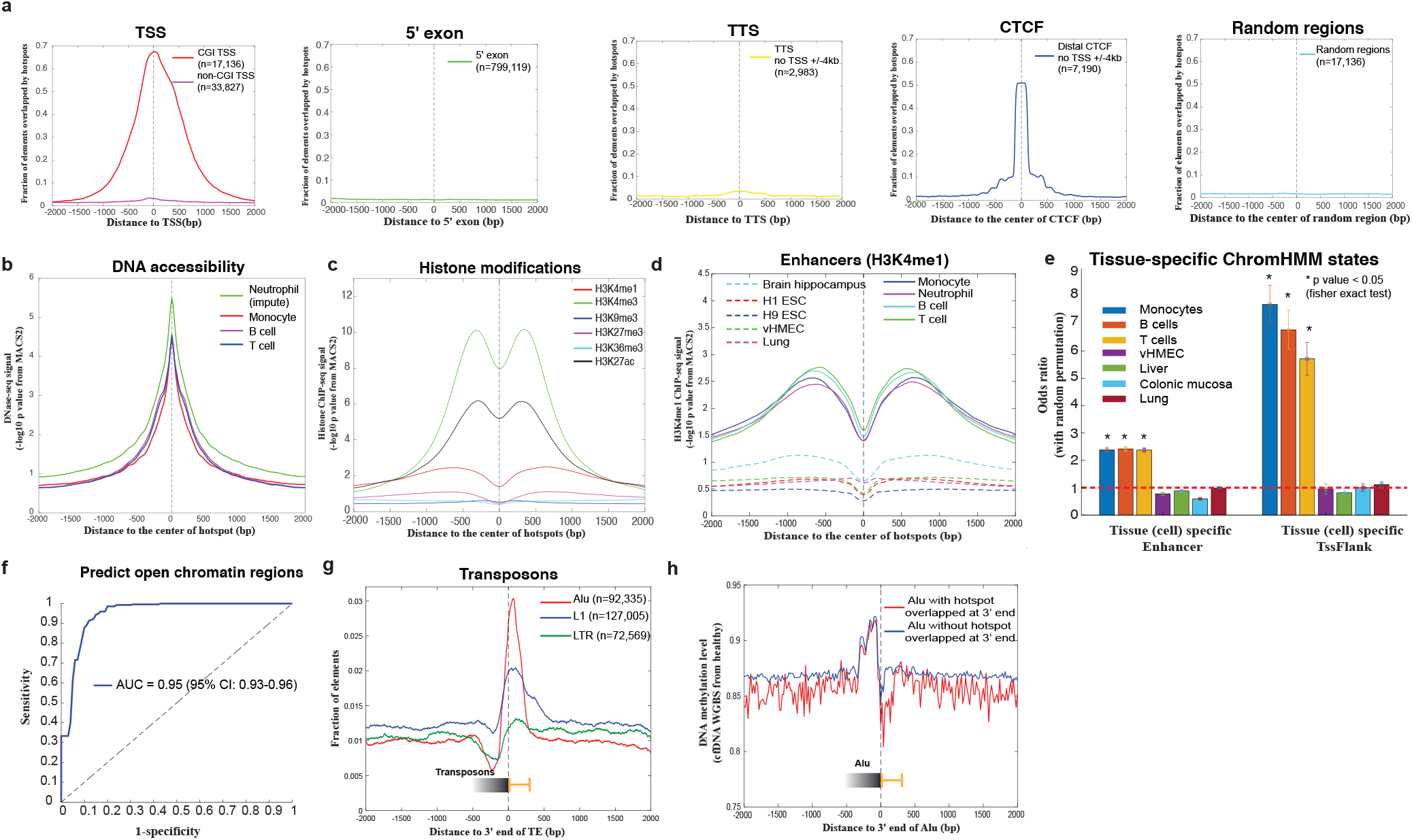
CfDNA fragmentation hotspots are enriched at active gene-regulatory regions in healthy. (a). The overlap of cfDNA fragmentation hotspots (BH01, healthy) and CGI Transcription Starting Sites (TSSs), non-CGI TSSs, 5’exon boundary (no TSS and CTCF within +/-2 kb), Transcription Termination Sites (TTSs)(no TSS and CTCF within +/-2 kb), CTCF transcription factor binding sites (no TSS within +/-4 kb), and random genomics regions. (b) The DNA accessibility levels from hematopoietic cells around the cfDNA fragmentation hotspots(BH01, healthy). (c).The histone modification levels from monocytes around the cfDNA fragmentation hotspots(BH01, healthy). (d). The H3K4me1 histone modification levels from hematopoietic (solid lines) and non-hematopoietic (dashed lines) cells around the cfDNA fragmentation hotspots(BH01, healthy). (e). The enrichment of hotspots at tissue-specific chromHMM states (Enhancer and TssFlank). Odds ratio is compared with matched random regions (matched chromosome and length, repeated 10 times). Error bar is based on the 95% confidence interval. P value is calculated based on Fisher exact test. (f). ROC curve for the prediction of open chromatin regions by using cfDNA fragmentation level at the hotspots from the constitutively open and closed regions. (g). The overlap of cfDNA fragmentation hotspots (BH01, healthy) and 3’end of transposons (Alu, L1, and LTR). (h). The cfDNA methylation level from healthy individuals (Sun eta l. 2015 PNAS)[22] around the 3’end of Alu that overlapped or not overlapped with the cfDNA fragmentation hotspots (BH01, healthy).

We further asked if we could predict the open chromatin regions by using fragmentation alone. Neutrophils are one of the major contributors to the cfDNA in healthy individuals (20-60%)[21,22]. However, the open chromatin regions in neutrophils are still missing. Thus, we utilized the matched constitutively open regions and closed regions across different cell types to benchmark the accuracy that we can detect the open chromatin regions by the fragmentation level. We achieved the 0.95 (95% CI: 0.93-0.96) area under the curve (AUC) to predict the known open chromatin regions (Fig. 2f, Details in Methods), further suggesting the strong link between fragmentation hotspots and open chromatin regions.

We next asked if we could detect other unknown regulatory potentials from cfDNA fragmentation hotspots. We collected 523 publicly available open chromatin region datasets measured by DNase-seq or ATAC-seq across different cell types (Details in Table S1). These cell types are the major known contributors to cfDNA in healthy non-pregnant individuals, including liver and rest or activated immune cells from the Roadmap Epigenomics Consortium, ENCODE, BLUEPRINT, and other publications[15,23,25–27]. Interestingly, after excluding the potential overlap with all these known open chromatin regions, we noticed a high enrichment of hotspots not within but right after the 3’ end of transposable elements (TEs). To exclude the possible artifact of reads mapping caused by the sequence composition bias, we examined the distribution of mappability and G+C% content right after the 3’ end of TEs and did not notice the significant bias there (Fig. 2g, Fig. S5a, b). The motif enrichment results at these hotspots right after the 3’end of TEs further suggested the high enrichment of pioneer transcription factors, such as OCT (POU, Homeobox), which usually bind the nucleosome-occupied regions (Fig. S5c)[28]. Moreover, we observed the differences in DNA methylation at the same regions (right after the 3’end of Alu) with or without the overlap of hotspots, which indicates the potential functional association between hotspots and the local epigenetic status, besides nucleosome occupancy, after the 3’end of TEs (Fig. 2h).

Taken together, in healthy individuals, these *de novo* characterized cfDNA fragmentation hotspots are highly enriched in open chromatin regions and active gene regulatory elements and can potentially reveal other unknown regulatory elements from cfDNA WGS.

### Cell-free DNA fragmentation hotspots reveal the potential gene regulatory aberrations in early-stage cancer

We next sought to explore whether or not the cfDNA fragmentation dynamics in the hotspots can reflect the aberrations of gene regulatory elements in early-stage cancer. We collected the publicly available low-coverage cfDNA WGS (∼1X/sample) from 90 patients with early-stage hepatocellular carcinoma (HCC, 85 of them are Barcelona Clinic Liver Cancer stage A, 5 of them are stage B) and 32 healthy individuals from the same study[29,30]. Since these cfDNA WGS are all sequenced with low coverage, we estimated the minimum number of fragments required by CRAG (see Supplementary Methods and Fig. S6) and pooled these low coverage cfDNA WGS to obtain enough fragments for the hotspot calling in each condition. The unsupervised hierarchical clustering of the top 10,000 most variable hotspots showed a clear fragmentation dynamic between early-stage HCC and healthy (Fig. 3a, Fig. S7). The volcano plot of the false discovery rate (FDR, two-sample t-test) and z-score difference of IFS between HCC and healthy across all the fragmentation hotspots showed a large fraction of hypo-fragmented hotspots in early-stage HCC (Fig. 3b).

**Fig. 3.**
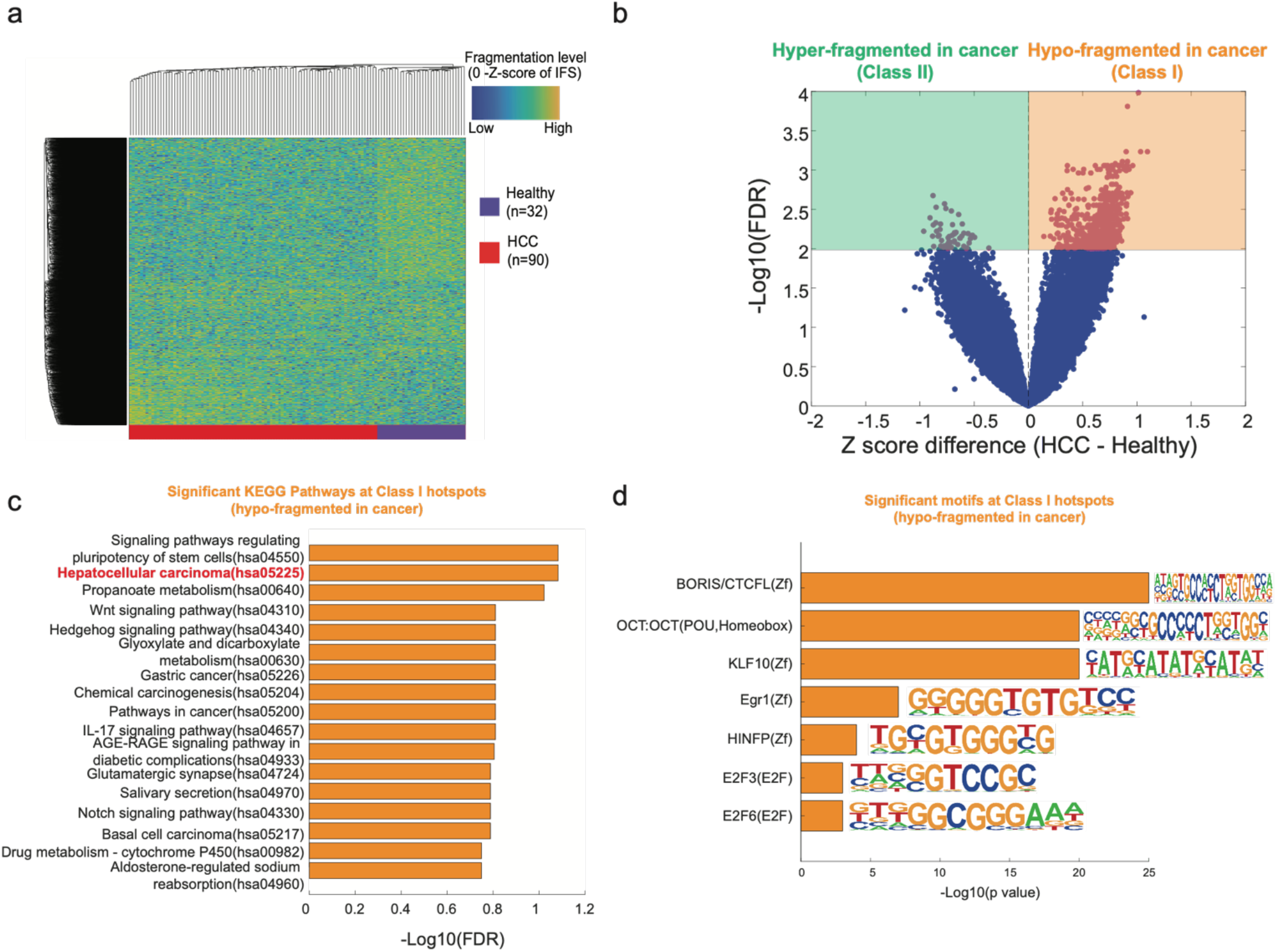
The aberrations of cfDNA fragmentation patterns at hotspots in early-stage cancers. (a). Unsupervised clustering on the Z-score of IFS at the top 10,000 most variable cfDNA fragmentation hotspots called from HCC and healthy samples. Hotspots are selected based on the variance of the IFS values across all the samples. (b). Volcano plot of z-score differences and -log10 FDR-value (two-way student’s t-test) for the aberration of IFS in cfDNA fragmentation hotspots between early-stage HCC and healthy. (c).The significant KEGG pathways for the genes associated with Class I hotspots. Cistrome-GO was utilized to associate hotspots with their targeted genes. (d). The motifs that are significantly enriched within Class I hotspots.

To further understand the molecular mechanism behind these fragmentation aberrations at the hotspots, we split the significantly differentiated fragmentation hotspots (FDR<0.01) into two groups: Class I (Hypo-fragmented in cancer) and Class II (Hyper-fragmented in cancer) (Fig. 3b, Table S2). We associated these hotspots with their targeted genes by CistromeGO and identified the enrichment of Gene Ontology Biological Processes (GO BPs) at these genes (Fig. S8)[31]. Genes associated with Class I hotspots are enriched in “cell adhesion” related GO BPs. For example, Epithelial Cell Adhesion Molecule (EpCAM) genes within the GO: 0098742 were considered the marker of HCC cancer stem cells for a long time[32,33]. Genes associated with Class I hotspots are relatively enriched in “cysteine endopeptidases”, “apoptosis”, and “purine biosynthesis” related GO BPs, which were all associated with cancer progression and invasion in the previous studies[34,35]. We also characterize the KEGG pathway enrichment at the Class I hotspots (Hypo-fragmented in cancer), which are highly associated with HCC initiations, such as Hepatocellular Carcinoma (hsa05225) and signal pathway regulating pluripotency of stem cells (hsa04550) (Fig. 3c). Interestingly, we found the motif enrichment of BORIS/CTCFL at the Class I hotspots (Hypo-fragmented in cancer) but not at the Class II hotspots (Hyper-fragmented in cancer) (Fig. 3d), which suggested the potential associations with the changes in three-dimensional chromatin organizations.

To understand the cell type specificity of these cancer-specific hotspots, we performed the enrichment analysis at chromatin states from different cell types. In the Epigenome Roadmap studies, Enhancer and TssFlank are considered to be mostly cell-type-specific[23]. Compared to Class I hotspots (Hypo-fragmented in HCC, i.e., open in healthy), we found that the Class II hotspots (Hyper-fragmented in HCC, i.e., open in HCC) are significantly enriched in cell-type-specific chromHMM states from liver and liver cancer (HepG2) but not other cell types (Fig. S9a).

In summary, in early-stage cancer, we found the global aberrations of fragmentation patterns at the cfDNA fragmentation hotspots, which bring together the signals mostly from peripheral immune cells and potentially small fractions from tumor tissues. These aberrations at the hotspots are highly associated with the alterations of regulatory elements and genes related to the initiation of cancer.

### Cell-free DNA fragmentation hotspots for the detection and localization of multiple early-stage cancers

Next, we asked if we could utilize the cfDNA fragmentation hotspots for the diagnosis of early-stage cancer. The diagnosis of early-stage HCC is usually compared with not only healthy individuals but also patients with liver diseases. Thus, we collected additional cfDNA WGS datasets in 67 patients with chronic HBV infection and 36 patients with HBV-associated liver cirrhosis from the same study as above [29]. Unsupervised hierarchical clustering at the most variable hotspots showed the clear dynamics of the fragmentation patterns among early-stage HCC, HBV, Cirrhosis, and healthy controls (Fig. S10-11). Utilizing the ten-fold cross-validation, we identified hotspots only in the samples pooled in the training dataset to avoid the information leak to the test dataset. We utilized the z-score transformed IFS from the cfDNA fragmentation hotspots as the features for the classification by a linear Support Vector Machine (SVM) approach (Details in Methods). Overall, for the comparison between HCC and healthy, we obtained 91% sensitivity at 100% specificity (96% sensitivity at 100% specificity after GC bias correction) (Table S3-4, Fig. S12). For the comparison between HCC and all other non-cancer controls, we obtained 83% sensitivity at 100% specificity (88% sensitivity at 100% specificity after GC bias correction). Both comparisons showed higher performance than other methods, especially fragmentation at known regulatory elements, from the same dataset with the same data split (Fig. S13, Table S5-6).

We further extended our study from early-stage HCC to multiple other cancer types. We collected publicly available low-coverage cfDNA WGS data (∼1X/sample) from 208 patients across seven different kinds of cancer (88% in stage I-III, colon, breast, lung, gastric, bile duct, ovary, and pancreatic cancer) and 215 healthy controls in the same study[2,30]. We applied a similar strategy as the HCC study above for the hotspot calling (pool the samples in the training dataset). Across seven different types of cancer and healthy conditions, the z-score transformed IFS signals in the most variable fragmentation hotspots showed clear cancer-specific fragmentation patterns in both t-SNE visualization and unsupervised hierarchical clustering (Fig. 4a-b, Fig. S14, Details in Supplementary Methods). We also performed the enrichment analysis at chromatin states similar to that in HCC above. Compared to Class I hotspots (Hypo-fragmented in the lung cancer, i.e., open in healthy), we found that the Class II hotspots (Hyper-fragmented in lung cancer, i.e., open in cancer) are significantly enriched in cell-type-specific chromHMM states from lung cancer (A549)(Fig. S9b).

**Fig. 4.**
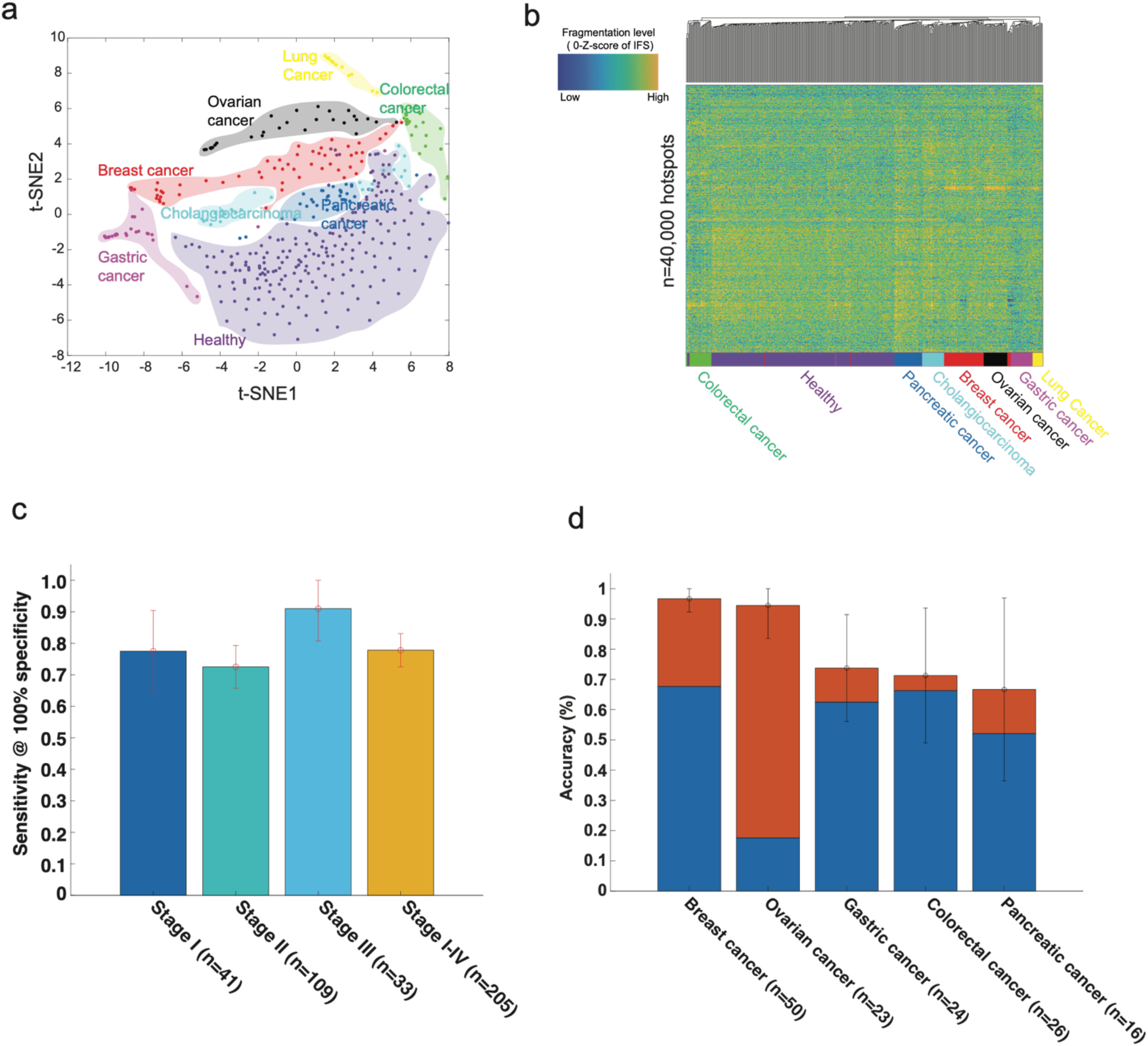
The detection and localization of multiple early-stage cancers. (a). t-SNE visualization on the Z-score of IFS (after GC bias correction) at the most variable cfDNA fragmentation hotspots (one-way ANOVA test with p-value < 0.01) across multiple different early-stage cancer types and healthy conditions. (b). Unsupervised clustering on Z-score of IFS (after GC bias correction) at the top 40,000 most variable cfDNA fragmentation hotspots across multiple different early-stage cancer types and healthy conditions. (c). The sensitivity across different cancer stages at 100% specificity to distinguish cancer and healthy condition by using IFS (after GC bias correction) at cfDNA fragmentation hotspots called at training dataset. Error bars represent 95% confidence intervals. (d). Percentages of patients that were correctly classified by one of the two most likely types (sum of orange and blue bars) or the most likely type (blue bar). Error bars represent 95% confidence intervals.

By 10-fold cross-validation, the linear SVM model showed a consistent high classification performance across different stages for its high sensitivity at high specificity (72% sensitivity to 91% sensitivity at 100% specificity, Fig. 4c, Table S7). Across different cancer types, we achieved 48% to 95% sensitivity at 100% specificity. Particularly, at 100% specificity, we achieved 95% sensitivity (95% CI: 85%-100%) in colorectal cancer, 93% sensitivity (95% CI: 85%-100%) in breast cancer, and 90% sensitivity (95% CI: 80%-100%) in gastric cancer, which of these are poorly detected at high specificity level by other liquid biopsy approaches in the same dataset [2,36–39]. (Fig. S15-16, Table S7, Table S8). In the other cancer types, the performance is largely comparable to the previous results [2]. Since the cfDNA cancer diagnosis is mostly affected by the tumor fractions, even in early-stage cancers, we estimated the tumor fractions in each sample by the CNV-based approach (ichorCNA)[40]. A recent study suggested that the increases in signal breadth could increase the sensitivity even with a low tumor burden in cfDNA[41]. *De novo* hotspot characterization will expand the signals and thus increase the sensitivity. Our approach indeed showed a consistently high performance across samples with different tumor fractions (Fig. S17).

To validate the model performance in an independent test dataset, we generated low-coverage cfDNA WGS at plasma from two types of cancers: early-stage HCC (n=8, stage I-II) and breast cancer (n=25, stage I-III), together with their matched healthy controls (1:1 matched age, gender, alcohol usage, and smoking history, meta-data details in Table S9). We utilized the model trained in Cristiano 2019 data (Breast vs. Healthy) and Jiang 2015 data (HCC vs. Healthy) directly on the test set. We obtained high performance in both independent datasets and showed superior performance to other approaches, especially known regulatory elements, in the same test sets (Fig. S18, Table S10). However, the ROC curve could show perfect separation between cancer and non-cancer controls in the test set but fail to validate either the sensitivity or the specificity based on the fixed cutoff when applying to the clinical cases. To further validate the clinical application of our approach, we performed the batch effect correction at the training and test set (Details in Supplementary Methods, Fig. S19). We fixed the cut-off of our model in the training set of the training dataset and compared the performance of the model at the validation set in the training dataset and the independent test set. Our model still showed high performance in the test dataset (87.5% sensitivity at 87.5% specificity in early-stage liver cancer and 56% sensitivity at 80% specificity in early-stage breast cancer) (Table S11).

Next, we asked whether or not we could identify the tissues-of-origin of cancer samples by using the fine-scale fragmentation levels alone. In the cancer positive samples identified above by the machine learning algorithm, without any clinical information about the patients, we further localized the sources of cancer to one or two anatomic sites in a mean of 80% of these patients across five different cancer types and 76% accuracy across six different cancer types. Furthermore, we were able to localize the source of the positive test to a single organ in a median of 62% of these patients (Fig. 4d, Table S12, Fig. S20) (Details in Methods). The prediction accuracy varies among tumor types, from 67% (95%CI: 36%-97%) in pancreatic cancer to 97% (95%CI: 92%-100%) in breast cancer (Fig. 4d and Table S12), but significantly higher than random choices by the frequency of samples in each cancer type (Fig. S21).

Overall, our proof-of-concept study on the publicly available cfDNA WGS dataset suggested that the *de novo* characterization of cfDNA fragmentation hotspots is a promising novel approach for the diagnosis and localization of multiple early-stage cancers.

## Discussion

In summary, we developed a computational approach, named CRAG, to *de novo* identify the cfDNA fragmentation hotspots by weighting fragment coverages with the fragment size information. The cfDNA fragmentation hotspots are highly enriched at open chromatin regions and active gene-regulatory elements. While in early-stage cancers, a significant proportion of these hotspots are hypo-fragmented. These hypo-fragmented hotspots in early-stage cancer are mostly enriched in GO terms and pathways related to the initiation of cancer, which further suggests the functional importance of these cancer-specific hypo-fragmented hotspots. In addition, the BORIS/CTCFL motif is enriched at these hypo-fragmented hotspots, which suggests the potential three-dimensional chromatin organization changes during the initiation of early-stage cancer that has been reported before but not revealed by the non-invasive cfDNA approaches [42]. Overall, our results suggested that the *de novo* characterization of fine-scale cfDNA fragmentation hotspots is critical to revealing the unknown gene-regulatory aberrations in pathological conditions.

Compared to the fragmentation studies at the known regulatory elements, such as TSS and TFBS, our *de novo* approach shows several advantages. First, *de novo* approach will expand the signal breadth. The tumor content in cfDNA is low in most early-stage cancers. Recent studies suggested that the increase of signal breadth could increase the sensitivity even with a low tumor burden in cfDNA[41]. The aberration of regulatory elements in cancer involves both genes and distal regulatory elements[14,43]. *De novo* characterization will expand the signals from known genes’ promoters to many distal regulatory elements and thus increase the sensitivity. Second, the landscape of known regulatory elements has not been well characterized in tumor and immune cells from early-stage cancer patients yet. The initiation of early-stage cancer involves the interaction between the tumor and the immune environment[44,45]. The chromatin accessibility landscape has recently been profiled in late-stage tumors in cancer patients and immune cells in healthy individuals[14,15]. However, the landscape of regulatory elements from early-stage tumors and the immune environment of patients with early-stage cancers are still not known. Therefore, it is challenging to utilize comprehensive regulatory element maps in tumor and immune cells for the study of early-stage cancers. Finally, a lot of distal regulatory elements do not contain well-defined motifs or TFBS. Moreover, tissue-specific and cancer-specific TFBS are also not well-defined across many diseases, especially early-stage cancers. Characterization of motifs and TFBS from sequence directly can not represent the whole aberration map of regulatory elements in early-stage cancers. Our benchmark results over the known regulatory elements also supported this conclusion (Fig. S18, Table S10).

The *in vivo* fragmentation process is complicated. There is a significant computational challenge to identify the fragmentation hotspots compared to the identification of fragmentation coldspots at nucleosome-occupied regions in cfDNA WGS[5]. For example, genomic regions with a higher fragmentation rate do not always indicate the open chromatin regions. Further, besides nucleosomes, both biological (e.g., DNA methylation and histone modifications)[46,47] and technical artifacts (e.g., G+C%, k-mer, and mappability)[17,48] can affect the measurements of fragmentation level. After excluding the known effects of open chromatin regions and technical artifacts, our genome-wide analysis here revealed the enrichment of hotspots after the 3’end of transposable elements and potentially associated with local DNA methylation level, which suggested the unknown origin of the cfDNA fragmentation processes.

Previous efforts had been made to characterize the nucleosome-free regions by using the depletion of coverages from MNase-seq/ChIP-seq assay [49]. The measurement of cfDNA fragmentation here, however, involves information from both fragment coverages and sizes. CRAG can be further improved by better integrating the fragment coverages and sizes, or even with more dimensions, such as the fragment orientation, jagged ends, and endpoint, to fully capture the spectrum of fragmentation. Also, G+C% bias is known to affect the peak calling result in ChIP-seq/ATAC-seq [50]. A better statistical model with the incorporation of GC normalization on both the fragment coverages and sizes will improve our method’s performance. PCR-free library preparation for WGS will also mitigate the concerns of GC bias and other sequencing artifacts [51].

Our study here on the detection and localization of early-stage cancer is still in the proof-of-concept stage. There are still several limitations. First, due to the limited number of publicly available early-stage cancer cfDNA datasets, the classification performance here is mainly evaluated by multi-fold cross-validation on a relatively small sample size cohort in each cancer type without strictly matched healthy controls, similar to other cfDNA WGS studies[2]. We only generated small-scale datasets from two cancer types for independent validations. Multiple independent large-scale prospective cohorts with strictly matched controls will be a better way to assess the power of our approach for the diagnosis of early-stage cancer. Second, previous studies suggested that pre-analytic differences in the patient populations could bring the artifact for fragmentomic studies and finally affect the diagnosis performance[52–57]. Unified sample collection, experimental workflow, and better computational approaches to adjust these cofounders are still needed. Third, we pooled the low-coverage WGS samples from the same condition for the hotspot calling, which may cause a problem with a small number of samples. Due to the random drop out of the fragment coverages and many genomic windows in the genome, the number of falsely discovered hotspots without any biological interpretations will increase. Our current strategy by filtering low mappability regions and correcting GC bias is helpful to reduce the false-positive rate for the hotspot detection. However, the accuracy of IFS signals at individual hotspots from each sample is still severely affected by the low coverage data. Recent efforts showed the possibility to integrate genome-wide mutational patterns at low-coverage WGS to enable the ultra-sensitive detection of cancer samples with low tumor burden[41], which is similar to our strategy for the IFS signals at low-coverage samples here. Since we narrow down the regions of interest, even with missing values at part of the loci, many other hotspots from the same sample will still provide informative signals rather than noises for the model to make the classifications. In the future, appropriate statistical models for the imputation of missing fragmentation patterns are still needed to mitigate the missing data problem. Lastly, in some cancer types, our fine-scale study here showed complementary classification performance compared with that in the previous large-scale fragmentation study in the same dataset [2]. For example, our results on gastric, breast, and colorectal cancer outperformed previous large-scale fragmentation studies, while for bile duct, pancreatic, and lung cancer, the performance is reversed. Future combinations of the fragmentation patterns at multi-scales and information from other modalities or clinical meta-data may further improve the performance.

## Materials and Methods

### Public Datasets

Public datasets used in this study are listed in Table S1.

### Preprocess of whole-genome sequencing data

The adapter was trimmed by Trimmomatic (v0.36)[58] in paired-end mode with the following parameters: ILLUMINACLIP:TruSeq3-PE.fa:2:30:10:2:true MINLEN:36. After adapter trimming, reads were aligned to the human genome (GRCh37, human_g1k_v37.fa) using BWA-MEM 0.7.15 [59] with default parameters. PCR-duplicate fragments were removed by samblaster (v0.1.24) [60]. Only high-quality autosomal reads were used for all downstream analyses (both ends uniquely mapped, either end with a mapping quality score of 30 or greater, properly paired, not supplementary alignment, and not a PCR duplicate). In addition, fragments shorter than 50bp and longer than 1000bp are excluded from downstream analysis.

### Preprocess of whole-genome bisulfite sequencing data

DNA methylation levels measured by WGBS in cfDNA were obtained from the previous publications (Details in Table S1) [22,61]. Single-end WGBS from cfDNA was processed by the following internal pipeline. Based on FastQC results on the distribution of four nucleotides along the sequencing cycle, the adapter was trimmed by Trim Galore! (v0.6.0) with cutadapt (v2.1.0) and with parameters “--clip_R1 10” and “--clip_R1 10 --three_prime_clip_R1 13”. After the adapter trimming, reads were aligned to the human genome (GRCh37, human_g1k_v37.fa) by Biscuit (v0.3.10.20190402) with default parameters. PCR-duplicate reads were marked by samtools (v1.9)[62]. Only high-quality reads were used for all the downstream analyses (uniquely mapped, mapping quality score of 30 or greater, and not a PCR duplicate). The methylation level at each CpG was called by Bis-SNP (v0.90) with default parameters in bissnp_easy_usage.pl [63].

### Identification of cfDNA fragmentation hotspots by CRAG

Fragment coverages and sizes are both essential parts of the cfDNA fragmentation patterns. However, popular peak calling tools, such as MACS2[64], cannot utilize the signals from two dimensions. Thus, we created an integrated fragmentation score (IFS) by weighting the fragment coverage based on the ratio of average fragment size in the window *versus* that in the whole chromosome. We also found that hotspots called by IFS showed a better enrichment at regulatory regions than those called by fragment coverage alone. We utilized a 200bp sliding window with a 20 bp step to scan each chromosome (autosome only). In the *i*^*th*^ window:

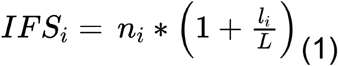

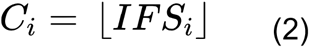

where *C*_*i*_ is the IFS score round down to the nearest integer in the *i*_*th*_ window, *n*_*i*_ is the number of fragments whose mid-points are located within the *i*_*th*_ window, *l*_*i*_ is the average fragment size in the *i*_*th*_ window, *L* is the average fragment size in the whole chromosome. Windows overlapped with dark regions or with average mappability scores smaller than 0.9 were removed. Dark regions were defined by the merged DAC blacklist and Duke Excluded from the UCSC Table Browser. Mappability score was generated by the GEM mappability program on the human reference genome (GRCh37, human_g1k_v37.fa, 51mer) [65].

The negative binomial (NB) model was previously utilized for the ChIP-seq peak calling[66]. Across the genome, the background IFS score is far from constant. The Poisson distribution, which requires a single fixed distribution mean (lambda), is not an ideal model to fit the overdispersed data. Instead, NB distribution can be viewed as a compound Poisson distribution, i.e., a continuous mixture of Poisson distributions of dispersed, Gamma-distributed lambdas. Thus, we found that the negative binomial model is better than the Poisson model by allowing the background IFS scores to vary across the genome. Here, we assumed the background *C*_*i*_ following the NB distribution.

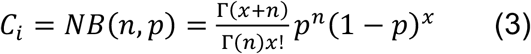

We denoted the sample mean and sample variance as *μ* and *ν*. Thus, we can estimate NB parameters as follows:

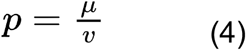

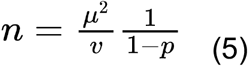

We utilized the NB model to test whether the *C*_*i*_ in the *i*_*th*_ window was significantly smaller than the local background (50 kb) and global background (the whole chromosome). In R (v4.2.0), we can calculate p-values using the following function:

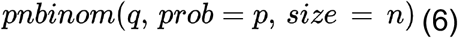

where *q* is the observed IFS score in the window. Based on Median Absolute Deviation (MAD), we identified the outlier of the IFS score (MAD > 5) and excluded them from the p-value calculation. For each window, we took the larger value between the local background and the global background p-value and then performed multiple hypothesis correction (Benjamini and Hochberg method). Only windows with an adjusted *p*-value smaller than a cut-off (FDR <= 0.2) were kept for further analysis. Finally, significant windows with a distance of less than 200bp to each other were merged as the final hotspots.

To remove the possible sequence composition bias caused by G+C% content, similar to the previous study[2], locally weighted smoothing linear regression (loess, span = 0.75) was utilized to regress out the GC covariates from the raw IFS score in each window. In R, we used the *loess* function for the calculation. The mean IFS score in each chromosome was added back to the residual value after the correction. The hotspots were called based on the corrected IFS finally.

To check the possible fragmentation bias caused by k-mer, we first calculated the expected IFS by using the average IFS at each possible type of dimer (16 types) across the genome. Then at each location, the adjusted IFS was calculated by dividing the original IFS with the expected IFS based on the dimer composition at that location. Finally, the adjusted IFS at each location was multiplied by the ratios between the average adjusted IFS and the average expected IFS in the same chromosome.

### Utilizing cfDNA fragmentation level to predict the open chromatin regions

We utilized the cfDNA fragmentation from healthy (BH01) as the benchmark dataset. Open chromatin regions from neutrophils, which release 20-60% cfDNA in healthy, are not available yet. We can not determine whether the ones that are predicted as open regions but not in the ATAC-seq/DNase-seq in immune cell types are false positive or still true positive but just the open regions in neutrophil, which is missing. Therefore, we used the conserved active chromatin regions and closed chromatin regions to benchmark the performance. We generated a balanced positive and negative group randomly sampled from two types of regions: (1) constitutively open regions: we used the -150 bp to 50 bp regions around the transcription start sites which are overlapped with TssA chromHMM states shared across all cell types (15-states chromHMM segmentation from NIH Epigenome Roadmap Consortium); (2) constitutively closed regions: we used the Quies chromHMM states shared across all cell types. We further randomly sampled the intervals from these constitutively closed regions with matched GC content and mappability as the constitutively open regions. We utilized the IFS score and k-mer (k=2) composition within these two types of regions as the features and applied the random forest model with default parameters (tree number = 100) in the setting of ten-fold cross-validation.

### Cancer early detection by cfDNA fragmentation hotspots

Here, we took the classification of liver cancer *vs*. healthy controls as an example. Ten-fold cross-validation was applied to evaluate the performance. In the training dataset, all the liver cancer samples and healthy samples were pooled to identify the hotspots, respectively. We used all the hotspots as the feature for the classification. It is well known that the sequencing depths will largely affect the number of peaks called in ChIP-seq and ATAC-seq[23]. In Cristiano et al. dataset [2], the sample size in the healthy group is ten times larger than that in any cancer type, which will lead to the uneven sequencing depths between healthy controls and cancers. Thus, by following the similar procedures in the previous publication[23], we downsampled the number of healthy controls to the same size as cancer before hotspot calling in each comparison (e.g., Breast cancer vs. Healthy). IFS before and after GC bias correction were both tested. IFS after GC bias correction was shown in the main figure for the classification. Only genomic regions at +/-100bp of the hotspot center were used to retrieve the IFS in each sample (the same strategy was used in PCA and unsupervised clustering analysis). The IFS at each corresponding hotspot was z-score transformed based on the mean and standard deviation at each chromosome of each sample. Finally, a support vector machine (SVM) classifier with linear kernel and default parameters (*fitcsvm* function at Matlab 2019b) was applied. At the testing dataset, the z-score transformed IFS in each sample was retrieved at the hotspot regions identified from the training set in that particular fold. The average AUC and 95% Confidence Interval (95% CI) of the AUC were calculated based on the classification results of the testing dataset across the ten folds. Specifically, we first compute the standard error of AUC (standard deviation divided by the square root of the iteration number). Then we multiply the standard error value by the z-score to obtain the margin of error. Finally, we add or subtract the margin of error from the mean AUC to obtain the confidence interval. To avoid the randomness of the data split, we repeated the cross-validation randomly 10 times.

### Tissues-of-origin predictions by cfDNA fragmentation hotspots

Only samples predicted as cancers were kept for the tissues-of-origin analysis. The saturation analysis of the fragment number needed for hotspot calling suggested that 200 million fragments are required to achieve the saturated performance (Fig. S6, Details in Supplementary Methods). Thus, pathological conditions with less than 200 million fragments in total were not used for the tissues-of-origin analysis (e.g., lung cancer). Bile duct cancer was at the boundary condition. Therefore, we performed the analysis with or without bile duct cancer. By 10-fold cross-validation, similar to that in the cancer early detection part, hotspots for each cancer type in the training set were identified. The z-score transformed IFS after GC bias correction in each sample was obtained as the feature. Since the total number of fragments in breast cancer is much larger than that in the other cancer types, we downsampled breast cancer to the median sample size across all the cancer types. The centroid in each cancer type was then calculated by the z-score transformed IFS across all the hotspots in the training set. In the testing dataset, each sample was assigned to the top two candidate cancers based on their distance to the centroids in each cancer type identified at the training set. The distance was calculated by *corr* function with ‘Type’ of ‘Spearman’ at Matlab 2019b. To further narrow down the best candidate cancer type, decision tree models (*fitctree* function at Matlab 2019b) were learned to identify the better candidate by all the hotspots in each possible pair of cancer types at the training set. Finally, we applied the corresponding decision tree model on the top two candidates to further characterize the best candidate at the testing dataset.

### Low coverage cfDNA WGS on plasma samples for independent validation

De-identified blood samples were collected under IRB approved studies 2012-3923 and 2012-3745 approved by the University of Cincinnati Institutional Review Board. Written informed consent was obtained from study participants to permit the collection and dispensing of de-identified biospecimens and associated clinical data for research. Plasma samples were provided by the UC Biorepository and UC Fernald Community Cohort. Clinical information about patients is provided in Table S9. CfDNA was isolated using the MagMAX Cell-Free DNA Isolation Kit (Applied Biosystems). The concentration and size distribution of cfDNA were measured by Qubit (Invitrogen) and BioAnalyzer (Agilent), respectively. Samples were randomly distributed into different batches for library preparation and sequencing. Case and their matched controls were always put into the same batch. Library construction was performed on 1ng of cfDNA using the KAPA HyperPrep Kit (Roche) and NEXTFLEX Unique Dual Index Barcodes (300nM final concentration, PerkinElmer). Libraries were sequenced on Illumina NovaSeq 6000 in PE150 mode.

## Acknowledgments

The authors greatly acknowledge Dr. Yuk Ming Dennis Lo and his circulating nucleic acids research group in the Chinese University of Hong Kong, Dr. Jay Shendure and his research group in the University of Washington, and Drs. Robert B. Scharpf, Victor E. Velculescu and their research group in the Johns Hopkins University School of Medicine for their cfDNA data. The authors acknowledge Dr. Li Wang for her suggestions and comments on the manuscript. The authors also acknowledge Jeanette M. Buckholz for the significant effort she invested in locating the study subjects. This work is supported by the computational resources from Biomedical Informatics (BMI) high performance computing cluster in CCHMC. This work was supported by the CCHMC start-up grant, Trustee Award, CCTST mentored pilot translational award, CCHMC innovation grant, and R56HG012360 from NHGRI to Y.L. This work also used the Extreme Science and Engineering Discovery Environment (XSEDE), which is supported by the National Science Foundation grant number ACI-1548562. This work used the XSEDE at the Pittsburgh Supercomputing Center (PSC) through allocation MCB190124P and MCB190006P. The authors acknowledge the Fernald infrastructure grant (R24ES028527) from NIEHS for the maintenance of the data and biospecimens, which were used as our validation samples.

## Author contributions

Y.L. conceived the study. Y.L. and X.Z. designed the methodological framework. X.Z. implemented the methods in MATLAB. H.Z. implemented the methods in R. X.Z., H.Z., and Y.L. performed the data analysis. H.F. generated the cfDNA WGS data for validation. K.L. D. M, S.M.P., and H.F. coordinated the plasma samples and provided input for the study design of the validation. Y.L., X.Z., H.Z, and H.F. wrote the manuscript together. All authors read and approved the final manuscript.

## Materials & Correspondence

For materials and correspondence, please contact Y.L

## Data and materials availability

The raw sequencing data (validation dataset generated by us) will be deposited at dbGap or EGA European Genome-Phenome Archive with controlled access (id will be available after acceptance). The de-identified fragment files mapped to hg19 are available at Zenodo.org (breast.tar and liver.tar, DOI: https://doi.org/10.5281/zenodo.6759611). All data needed to evaluate the conclusions in the paper are present in the paper and/or the Supplementary Materials. CRAG is implemented in MATLAB and R (v4.2.0). The source code is freely available on GitHub (https://github.com/epifluidlab/cragr.git) (R version) and (https://github.com/epifluidlab/CRAG.git) (MATLAB version) under the MIT license for academic researchers. The source code, readme files, hotspot location, intermediate analysis scripts, and intermediate results are available at Zenodo.org (DOI: https://doi.org/10.5281/zenodo.6759611).

## Ethics approval

This research study was approved by the Cincinnati Children’s Hospital Medical Center Institutional Review Board (#2019-0601) in accordance with the Declaration of Helsinki. De-identified plasma sample collection was approved by the University of Cincinnati Institutional Review Board (2012-3923 and 2012-3745).

