## Supplementary methods for "CRAG: *De novo* characterization of cell-free DNA fragmentation hotspots in plasma whole-genome sequencing"

*The saturation analysis of the fragment number required for CRAG.*

A group of fragmentation-positive and fragmentation-negative regions was generated for the benchmark. For fragmentation-positive regions, we chose the CGI TSS that are overlapped with conserved TssA chromHMM states (15-state chromHMM) shared across the cell types from NIH Epigenome Roadmap. Regions that are -50bp to +150bp around these active TSS were defined as the fragmentation-positive regions. For fragmentation-negative regions, we chose the same number of random genomic regions from conserved Quies chromHMM states shared across the cell types but with the same chromosome, region size, G+C% content, and mappability score as that in fragmentation-positive regions.

We downsampled the high-quality fragments in the BH01 dataset to 50 million fragments. We identified the hotspots at these downsampled datasets and calculated TP (true positive), FP (false positive), TN (true negative), FN (false negative) based on their overlaps with the benchmark regions generated above. F-score was calculated:

$$Fscore = 2 * \frac{\Pr ecision * \operatorname{Re} call}{\Pr ecision + \operatorname{Re} call} \quad (1)$$

in which, Precision and Recall were calculated using equation (2) and equation (3), respectively:

$$\Pr ecision = \frac{TP}{TP + FP} \quad (2)$$

$$\operatorname{Re} call = \frac{TP}{TP + FN} \quad (3)$$

The performance is saturated at ~0.9 F1-score with 200 million high-quality fragments. (Fig. S6).

*The enrichment analysis of the cfDNA fragmentation hotspots in gene-regulatory elements.*

The number of hotspots that overlapped with the regulatory element was counted by *bedtools v2* [1]. After filtering out the dark regions and low mappability regions (mappability less than 0.9), random genomic regions were generated with matched chromosomes and sizes. Fisher exact test (two-tail) was performed to calculate the enrichment of hotspots over the matched random regions.

*The Principal Component Analysis of the cfDNA fragmentation hotspots across different diseases.*

The cfDNA fragmentation hotspots were called at each pathological condition as described in the Methods. Principal Component Analysis (PCA) was performed on the z-score transformed IFS across all the fragmentation hotspots (*pca* function at Matlab 2019b).

*Unsupervised hierarchical clustering analysis of the cfDNA fragmentation hotspots across different diseases.*

The cfDNA fragmentation hotspots were called at each pathological condition as described in the Methods. Top *N* most variable hotspots were kept for the clustering (ranked by the variation across all the samples). Spearman's rank correlation was utilized to evaluate the distance among the samples. Also, weighted average distance (WPGMA, with 'weighted' as the parameter in *clustergram* function at Matlab 2019b) was applied together with the linkage method. In the Cristiano et al. dataset, one-way ANOVA (p-value < = 0.01) was applied to select the hotspots that showed the group-specific fragmentation patterns. Further, hotspots are ranked by the z-score difference between the samples within the group and outside the group. The top 5,000 hotspots in each group were finally visualized in the figure.

*The t-SNE visualization of the cfDNA fragmentation hotspots across different diseases.*

T-SNE (*tsne* function at Matlab 2019b) was utilized for the dimensionality reduction and visualization of the fragmentation dynamics in the hotspots across multiple cancer and healthy conditions. Hotspots with a p-value <= 0.01 (one-way ANOVA) were used for the analysis. Distance similarity was calculated by the Spearman correlation together with default parameters (*tsne* function at Matlab 2019b).

*The Gene Ontology and pathway analysis of the cfDNA fragmentation hotspots.*

Gene Ontology (GO) and KEGG pathway analysis of the cfDNA fragmentation hotspots was performed by Cistrome-GO[2].

*The motif analysis of the cfDNA fragmentation hotspots.*

Motif analysis of cfDNA fragmentation hotspots was performed by HOMER (v4.11) with the command ‘findMotifsGenome.pl hotspots_file hg19 output_file -size given’ [3]. Only motifs with a q-value of less than 0.01 were kept.

*The estimation of tumor fractions and copy number variations by ichorCNA.*

The ichorCNA v0.2.0[4] was run at 1Mb resolution with the normalization by the normal panel provided in the package together with G+C%, mappability, and the following parameters: --normal “c(0.75)” --ploidy “c(2)” --maxCN 5 --estimateScPrevalence FALSE --scStates “c(1,3)” --chrs “c(1:22)” .

*The batch effect correction and the performance evaluation at the independent test dataset.*

Principal Component Analysis (PCA) was performed on the z-score transformed IFS across all the fragmentation hotspots in public dataset and independent test dataset. There is indeed a batch effect between the training set (public data) and our independent test set (Fig. S19a). Therefore, we performed a batch effect correction. Specifically, we used *num.sv* and *sva* functions in *sva* package (R v4.2.0) to estimate the surrogate variables. *num.sv* suggests that there is only one major hidden surrogate variable. The underlying surrogate variable showed a high correlation with the data source and confirmed that the data source is indeed the only major contributor to the batch effect we observed (we also infer the gender by coverage ratio in chrX/chrY from the data). Finally, we apply the *ComBat* algorithm to correct the batch effects between the train and test datasets by their data source. After the correction, we randomly choose 2/3 of the samples in the training dataset (public data) to train the model (SVM classifier with linear kernel and default parameters). To balance the case and control categories, we down-sampled them to be equal during the training process. We evaluated the performance at the remaining 1/3 of the training set and chose the fixed cut-off to identify the positive and negative labels in the remaining 1/3 of the training set. We further applied this model and fixed the cut-off at the independent test set to identify the positive and negative labels. We repeated this process 100 times. The performance at 1/3 remaining data in the training set is shown in Table S11. The performance at the independent test set with the same cut-off and machine learning model is shown in Table S11. The high performance in the test set suggests that our method indeed has the potential clinical application.

#### Supplementary Figures

### Supplementary Figure 1

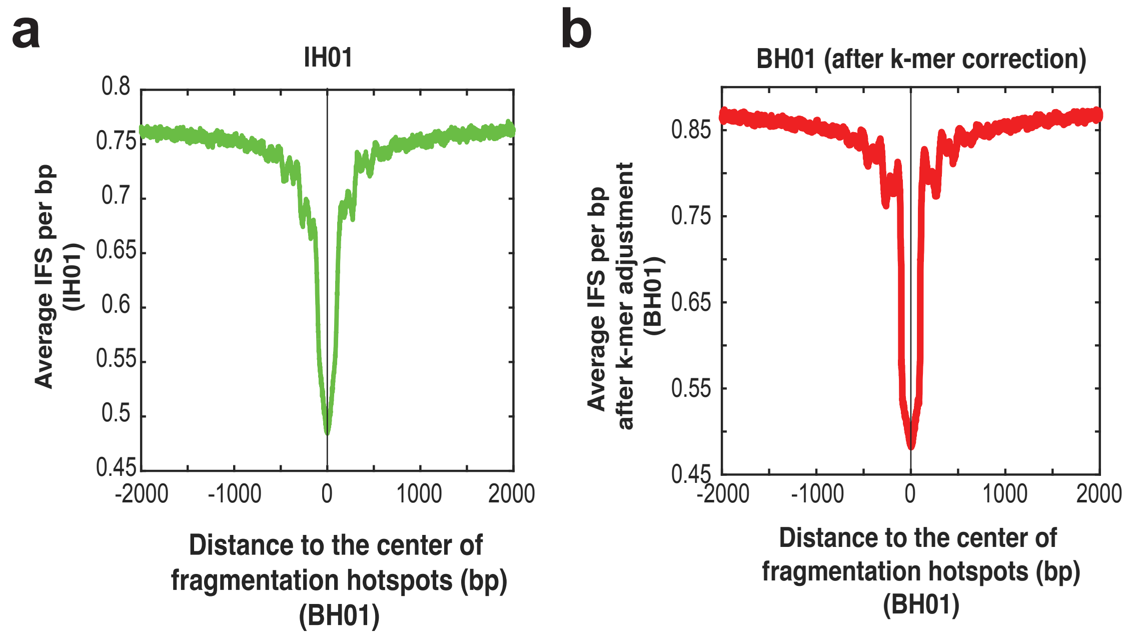


**Fig. S1. The fragmentation patterns near the cfDNA fragmentation hotspots**. The distribution of (a) IFS from IH01, and (b) adjusted IFS (after k-mer correction) from BH01 around the fragmentation hotspots called in the BH01 dataset.

#### Supplementary Figure 2

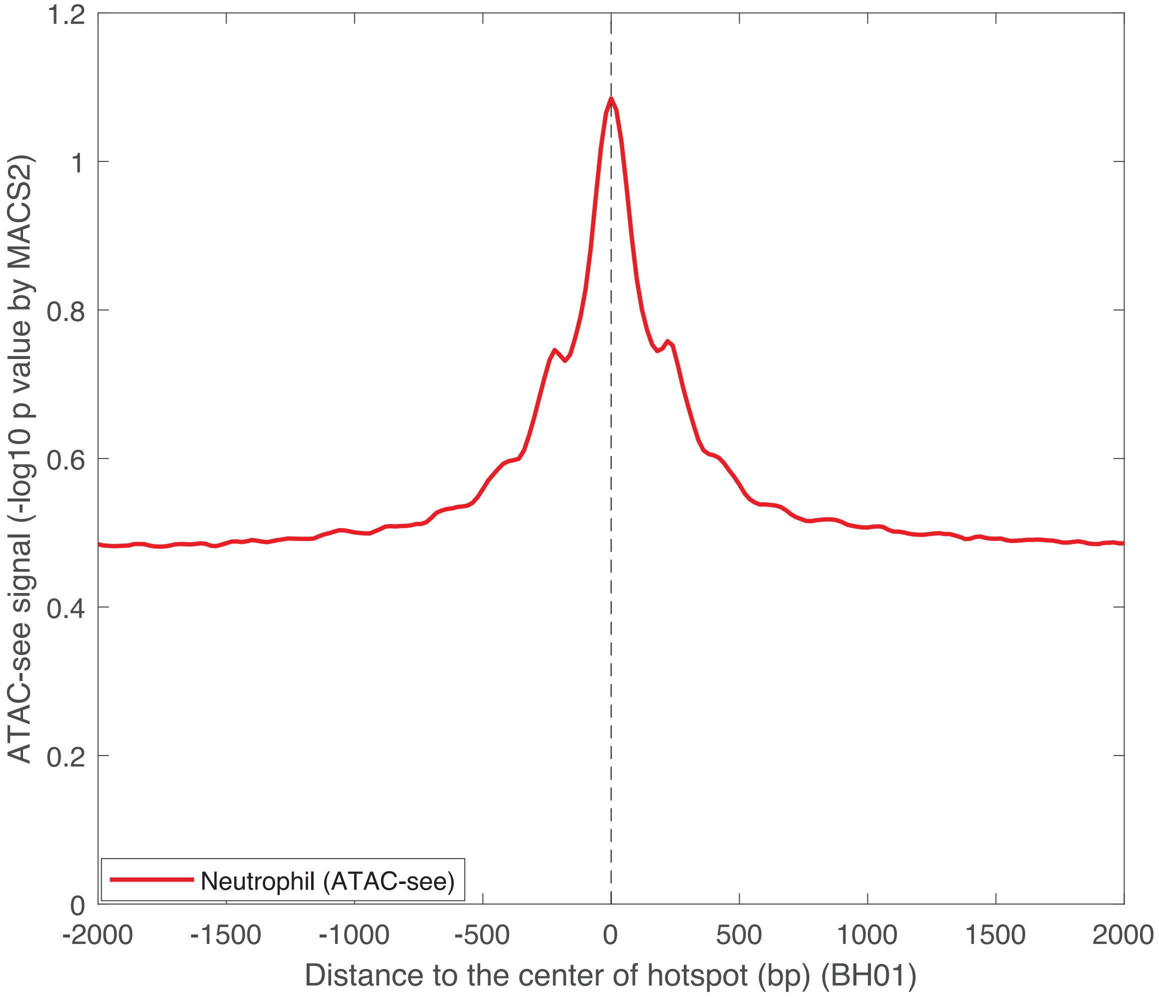


**Fig. S2. The enrichment of ATAC-see signals from neutrophils around the cfDNA fragmentation hotspots (BH01)**.

#### Supplementary Figure 3

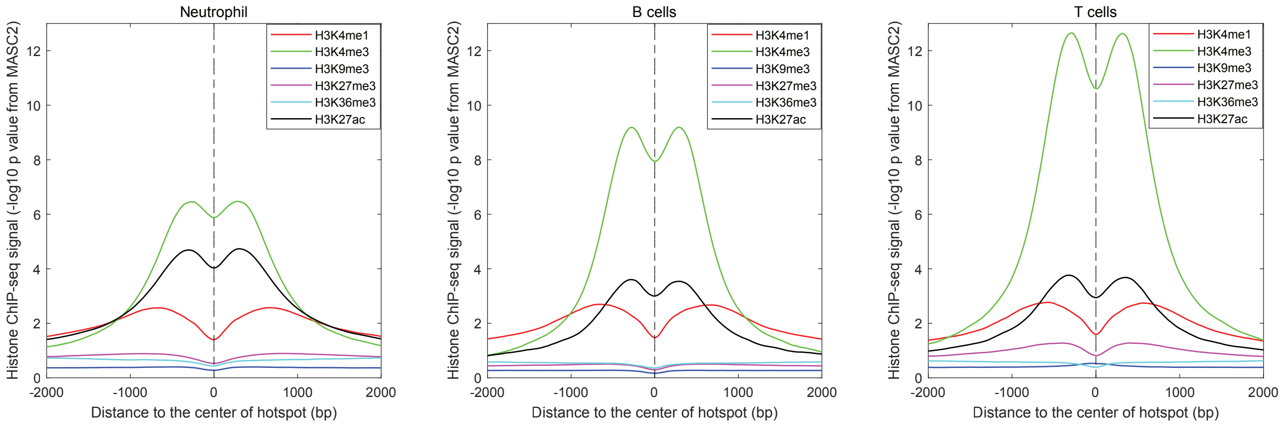


**Fig. S3. Epigenetic signals around cfDNA fragmentation hotspots (BH01).** The histone modification signal distributions (-log10 P-value calculated by MACS2, downloaded from Roadmap Epigenomics Consortium) from neutrophil, B cell, and T cell around cfDNA fragmentation hotspots (BH01).

**Supplementary Figure 4**

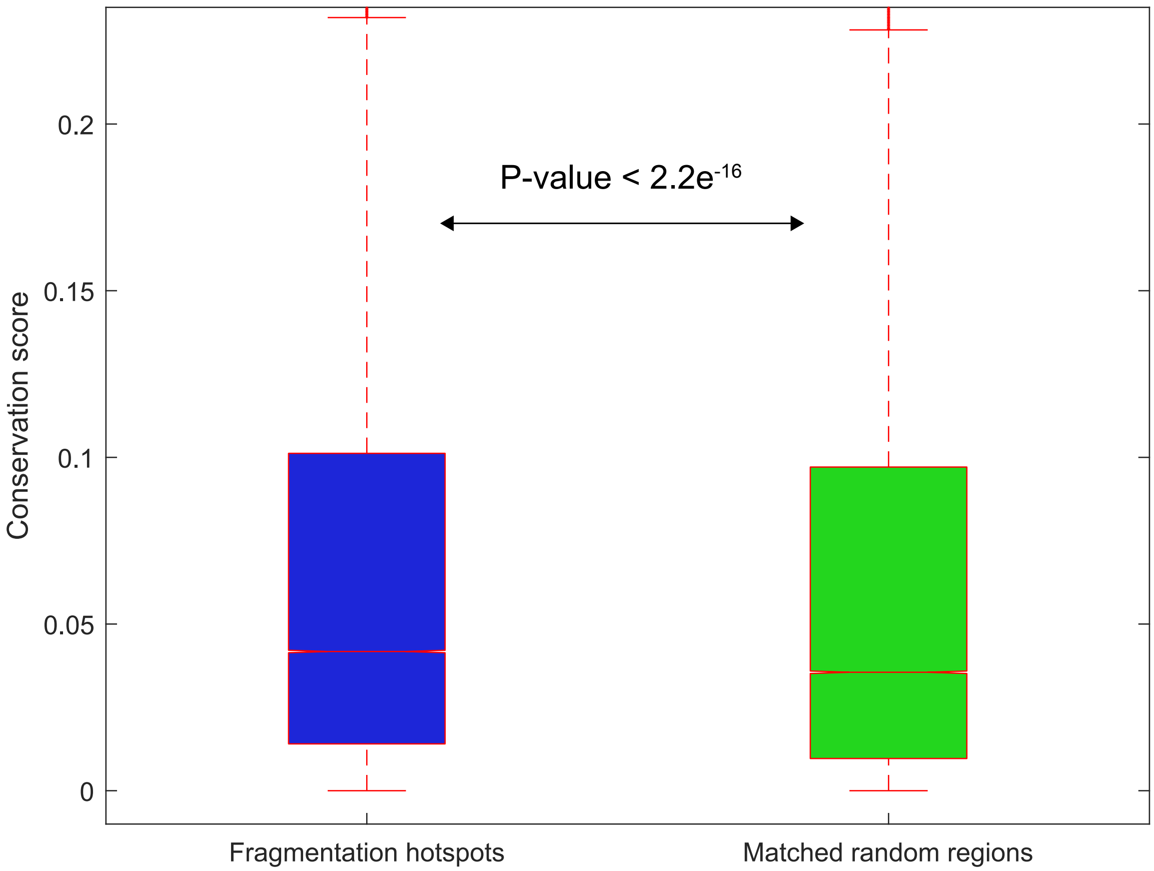


**Fig. S4. The boxplot of the conservation score (PhastCons) within cfDNA fragmentation hotspots and matched random regions.**

#### Supplementary Figure 5

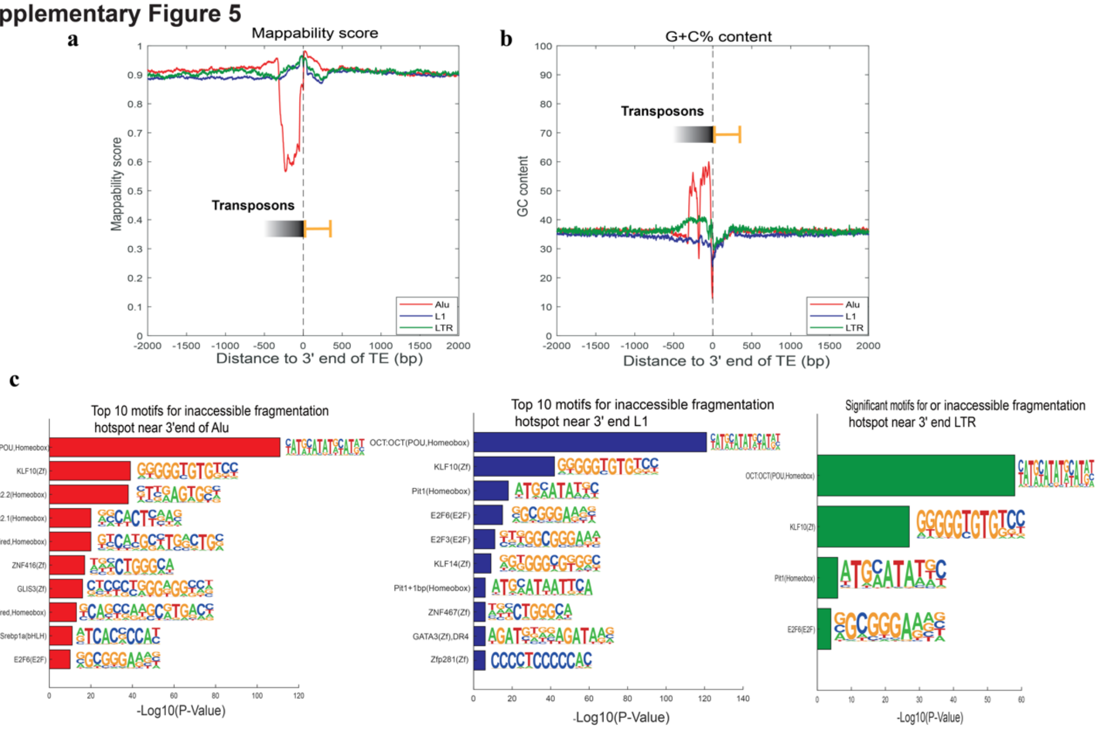

**Fig. S5. CfDNA fragmentation hotspots and transposable elements (TE)**. (a). The mappability score distribution at 3'end of TE. (b). The G+C% content distribution at 3'end of TE. (c). The top10 motif enrichment at hotspots after the 3'end of TE.

**Supplementary Figure 6**

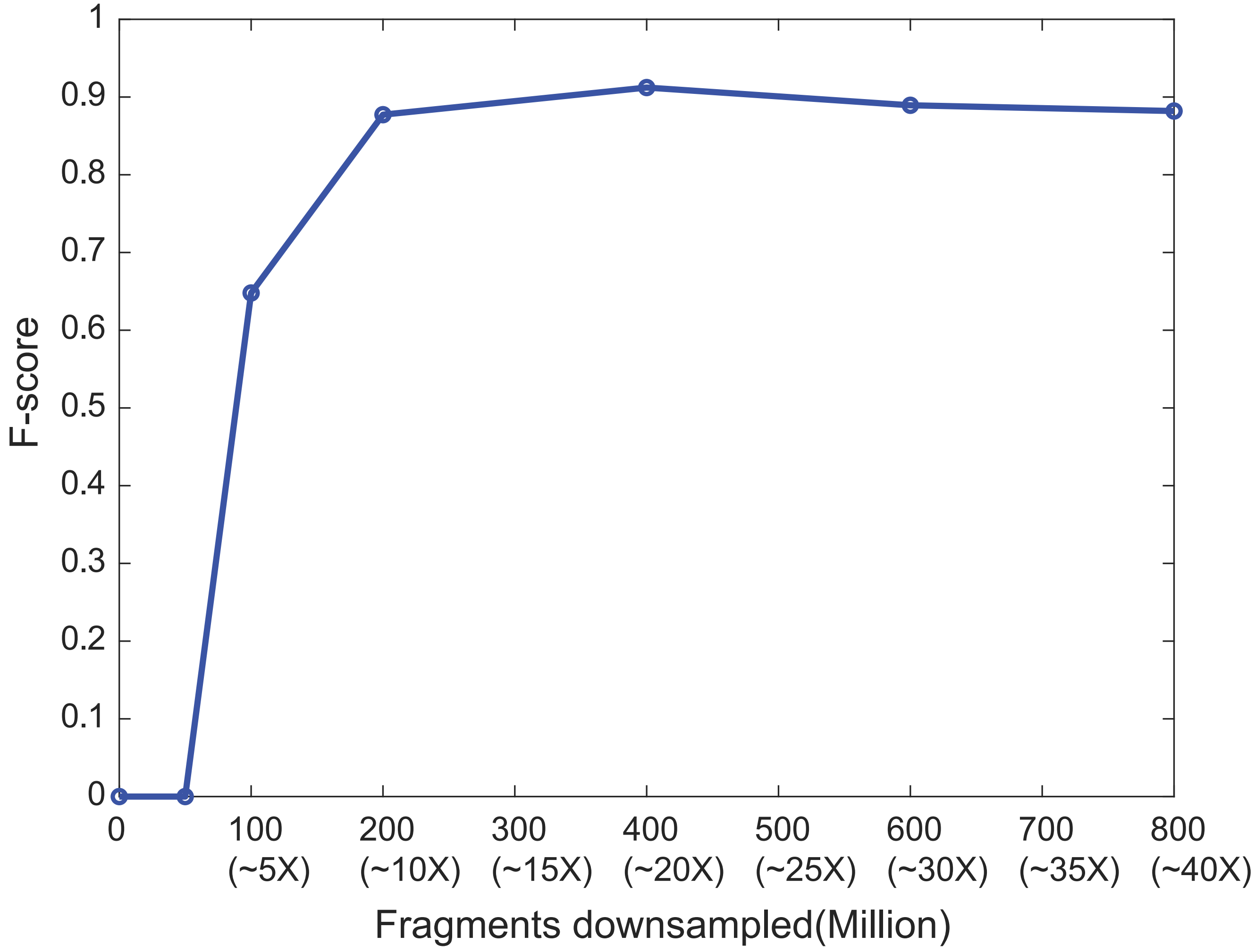


**Fig. S6. The power estimation for the cfDNA fragmentation hotspots called by CRAG with different numbers of fragments**.

### Supplementary Figure 7

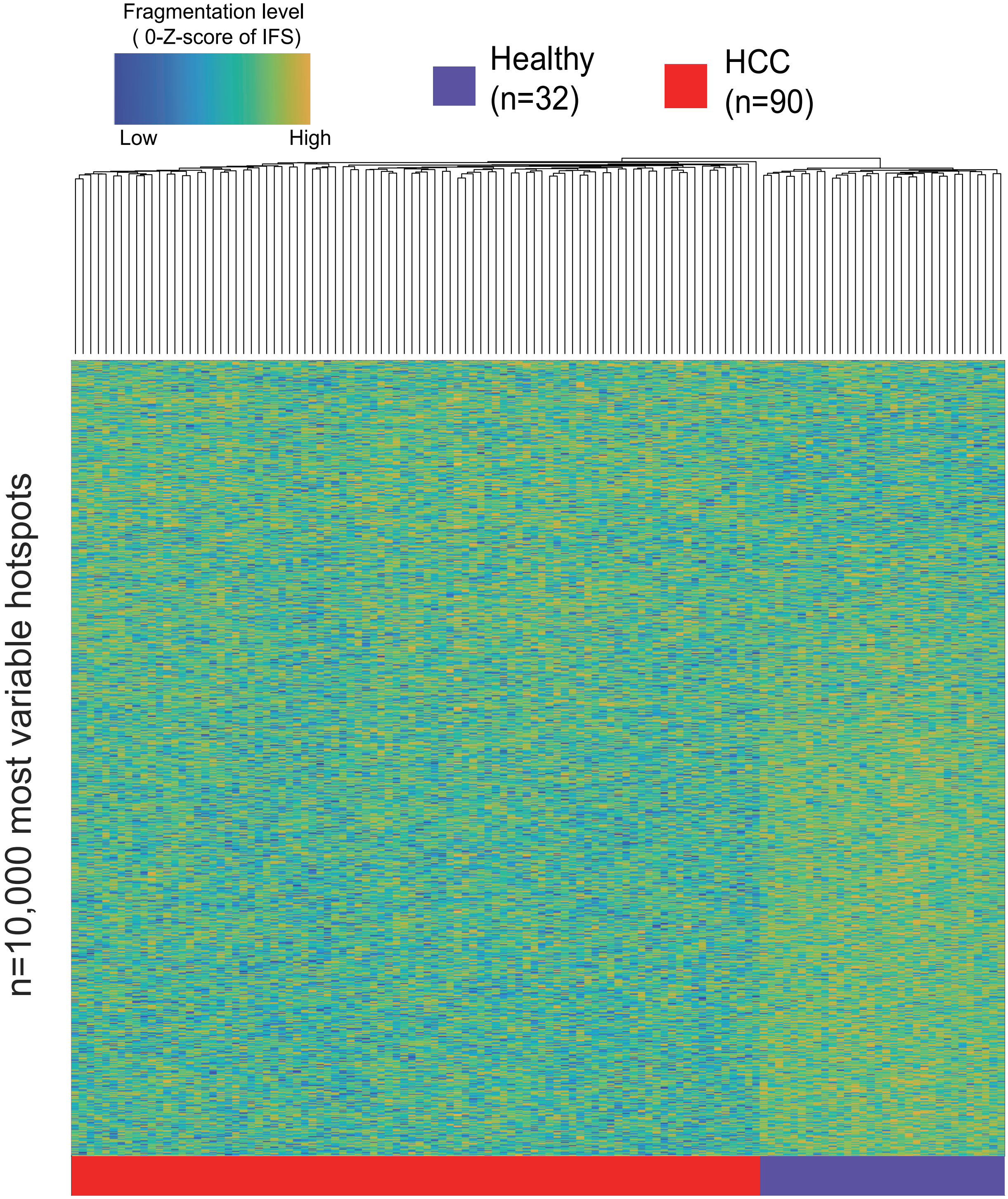


**Fig. S7. Unsupervised clustering on the Z-score of IFS at the top 10,000 most variable cfDNA fragmentation hotspots called from HCC and healthy samples (after GC bias correction).**

#### Supplementary Figure 8

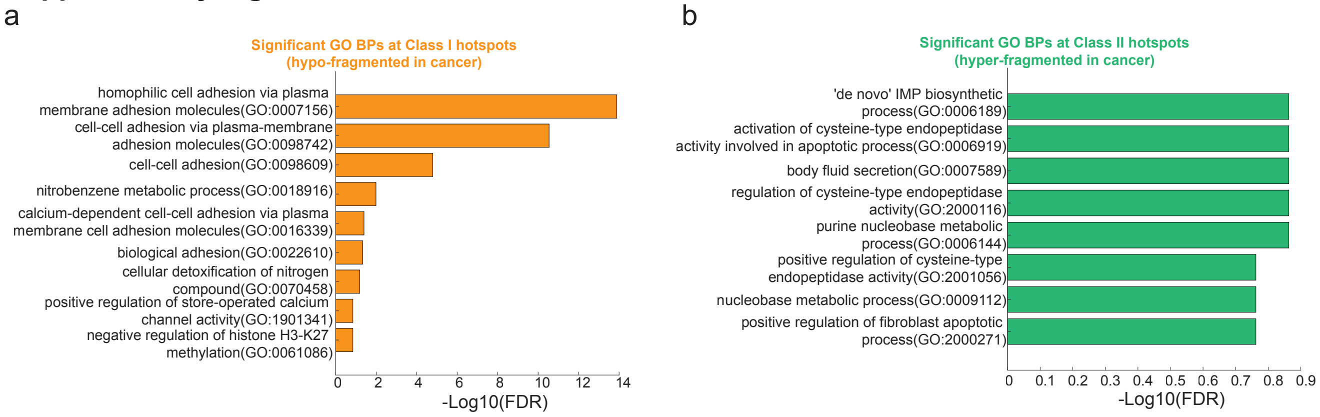


**Fig. S8. The significant GO BPs for the genes associated with (a) Class I and (b) Class II hotspots.**

Supplementary Figure 9

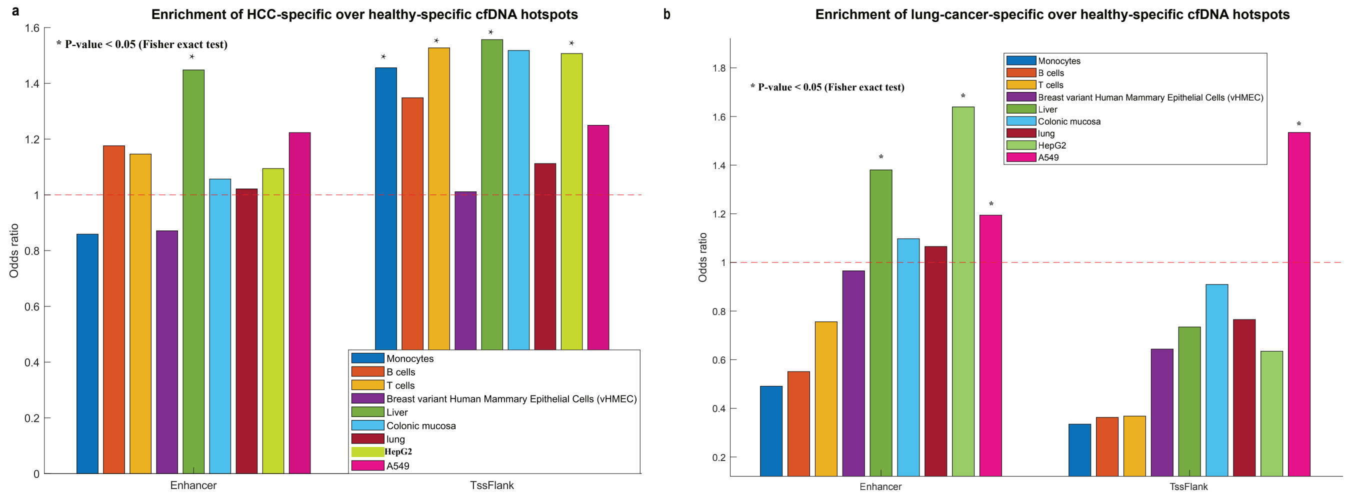


**Fig. S9. The enrichment of HCC-specific and lung-cancer-specific hotspots at different chromHMM states.** (a). The odds ratio is compared between HCC-specific (Class II) hotspots over healthy-specific (Class I) hotspots. P-value is calculated based on Fisher's exact test. (b). The Odds ratio is compared between lung-cancer-specific (Class II) hotspots over healthy-specific (Class I) hotspots. P-value is calculated based on Fisher’s exact test.

#### Supplementary Figure 10

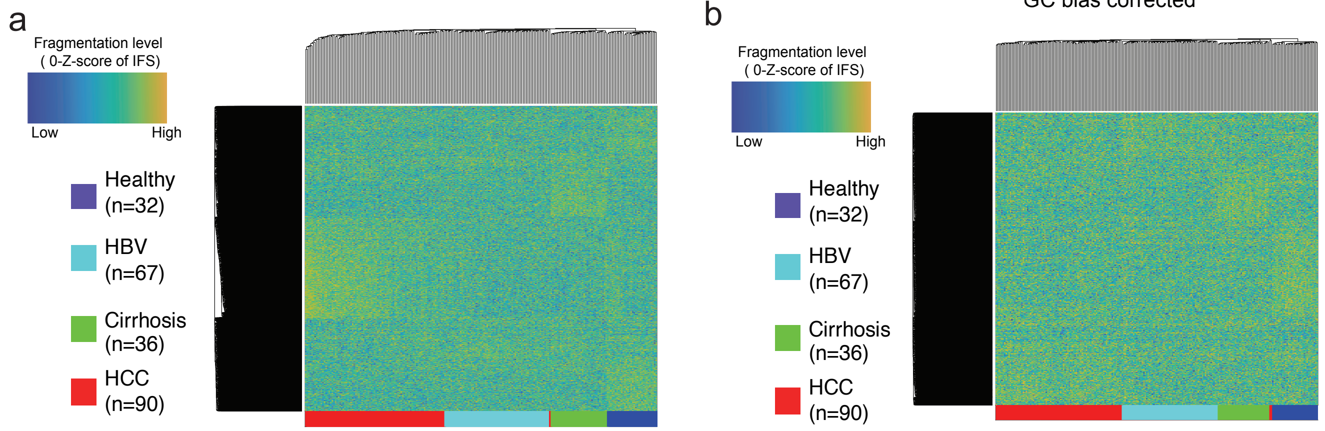


**Fig. S10. Unsupervised clustering on the Z-score of IFS at the top 10,000 most variable cfDNA fragmentation hotspots called from HCC (red), chronic HBV infection(cyan),HBV-associated liver cirrhosis(green), and Healthy(blue) samples** (a). Before and (b). After GC bias correction.

### Supplementary Figure 11

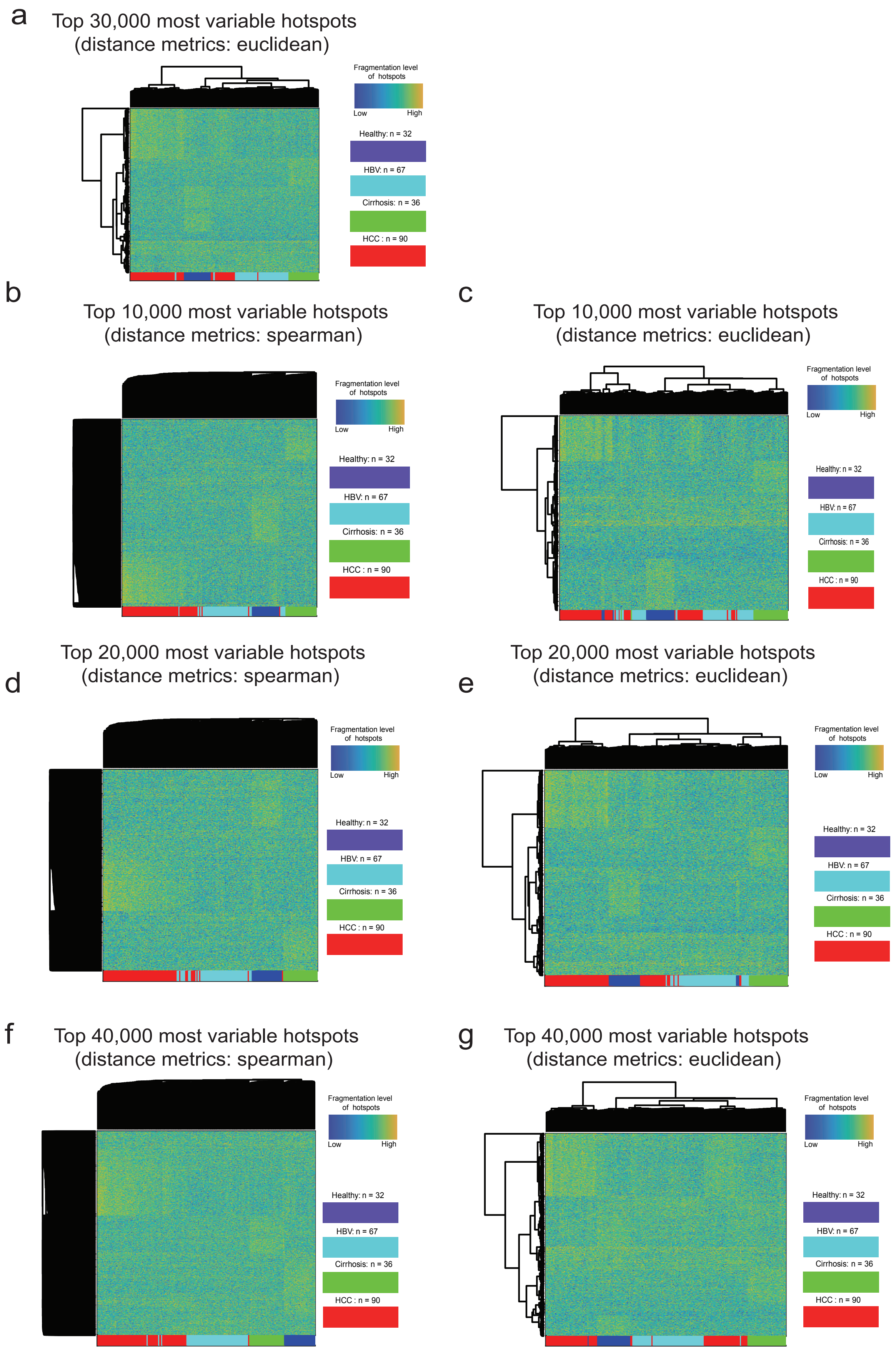

**Fig. S11. Unsupervised clustering on the Z-score of IFS at the most variable cfDNA fragmentation hotspots called from HCC, HBV-associated liver cirrhosis, chronic HBV infection, and healthy individuals**. (a). Clustering on the euclidean distance metrics from the top 30,000 most variable hotspots. (b). Clustering on the spearman correlation distance metrics from the top 10,000 most variable hotspots. (c). Clustering on the euclidean distance metrics from the top 10,000 most variable hotspots. (d). Clustering on the spearman correlation distance metrics from the top 20,000 most variable hotspots. (e). Clustering on the euclidean distance metrics from the top 20,000 most variable hotspots. (f). Clustering on the spearman correlation distance metrics from the top 40,000 most variable hotspots. (g). Clustering on the euclidean distance metrics from the top 40,000 most variable hotspots.

**Supplementary Figure 12**

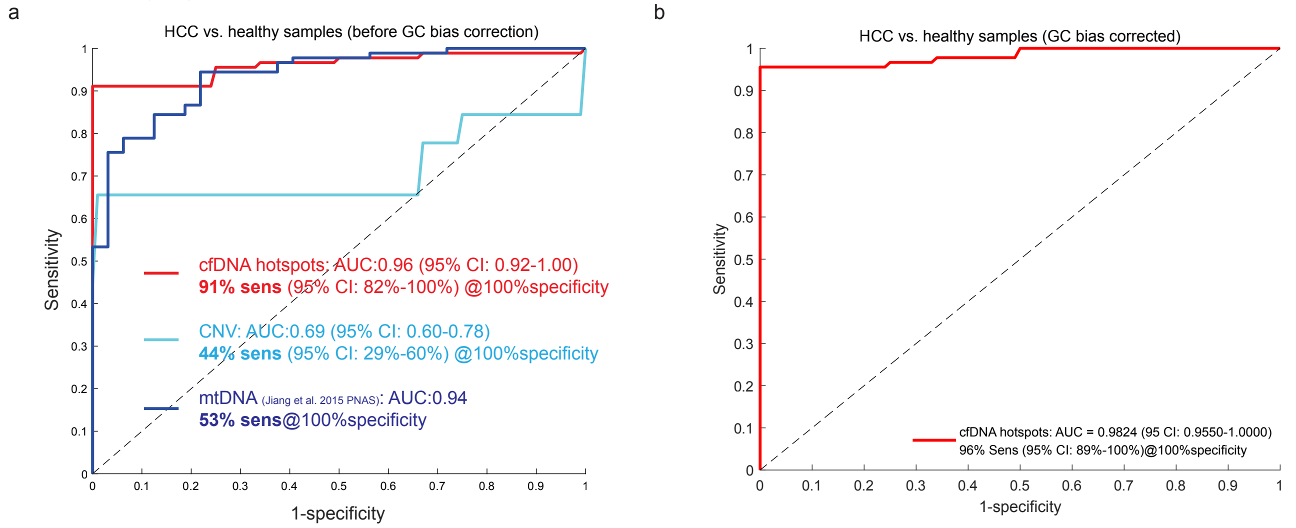


**Fig. S12. Receiver operator characteristics (ROC) for the detection of early-stage HCC.** IFS from cfDNA fragmentation hotspots (a) before and (b) after GC bias correction. The performance is compared with that by using copy number variations (cyan) and mitochondrial genome copy number analysis (blue).

**Supplementary Figure 13**

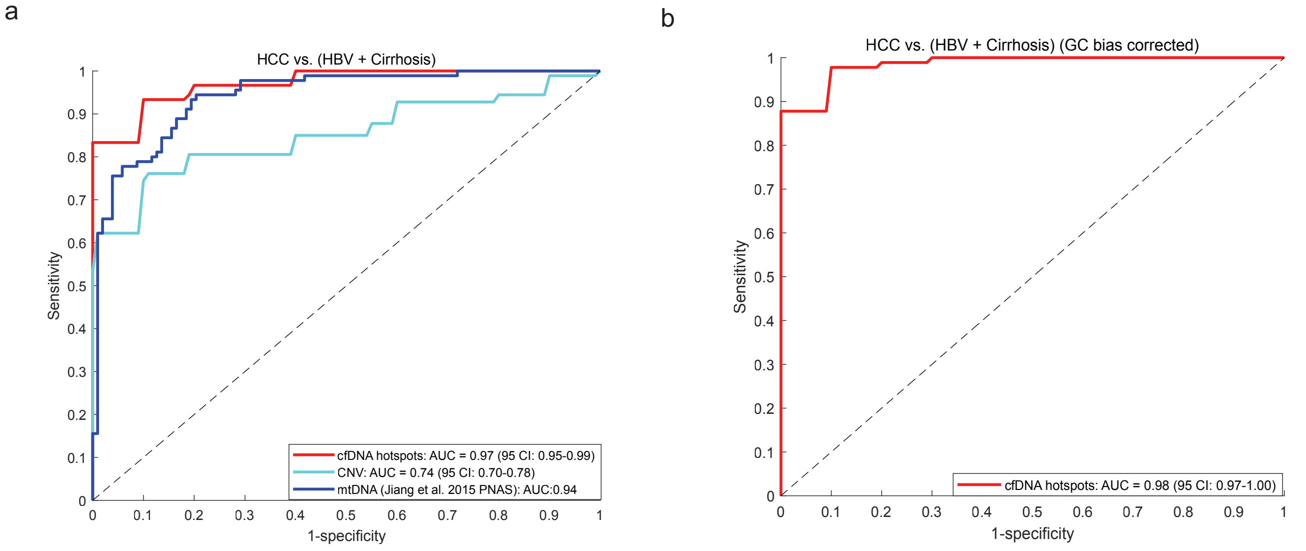


**Fig. S13. Receiver operator characteristics (ROC) to distinguish early-stage HCC with benign conditions (HBV-associated liver cirrhosis and chronic HBV infection) by using IFS from cfDNA fragmentation hotspots** (a). Before and (b). After GC bias correction.

**Supplementary Figure 14**

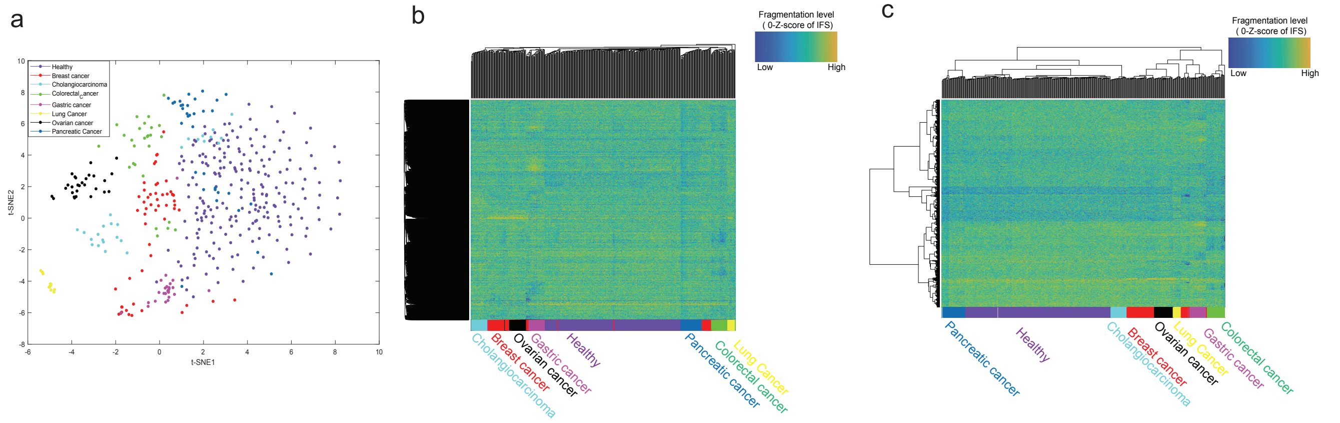


**Fig. S14. The aberrations of IFS (before GC bias correction) across multiple early-stage cancer and healthy**. (a). t-SNE visualization on the Z-score of IFS (before GC bias correction) at the top 40,000 most variable cfDNA fragmentation hotspots across multiple different early-stage cancer types and healthy. (b). Unsupervised clustering (WPGMA method on spearman correlation distance) on Z-score of IFS (before GC bias correction) at the top 40,000 most variable cfDNA fragmentation hotspots across multiple different early-stage cancer types and healthy. (c). Unsupervised clustering (Ward's method on euclidean distance) on Z-score of IFS (before GC bias correction) at the top 40,000 most variable cfDNA fragmentation hotspots across multiple different early-stage cancer types and healthy.

#### Supplementary Figure 15

a b c d e f g

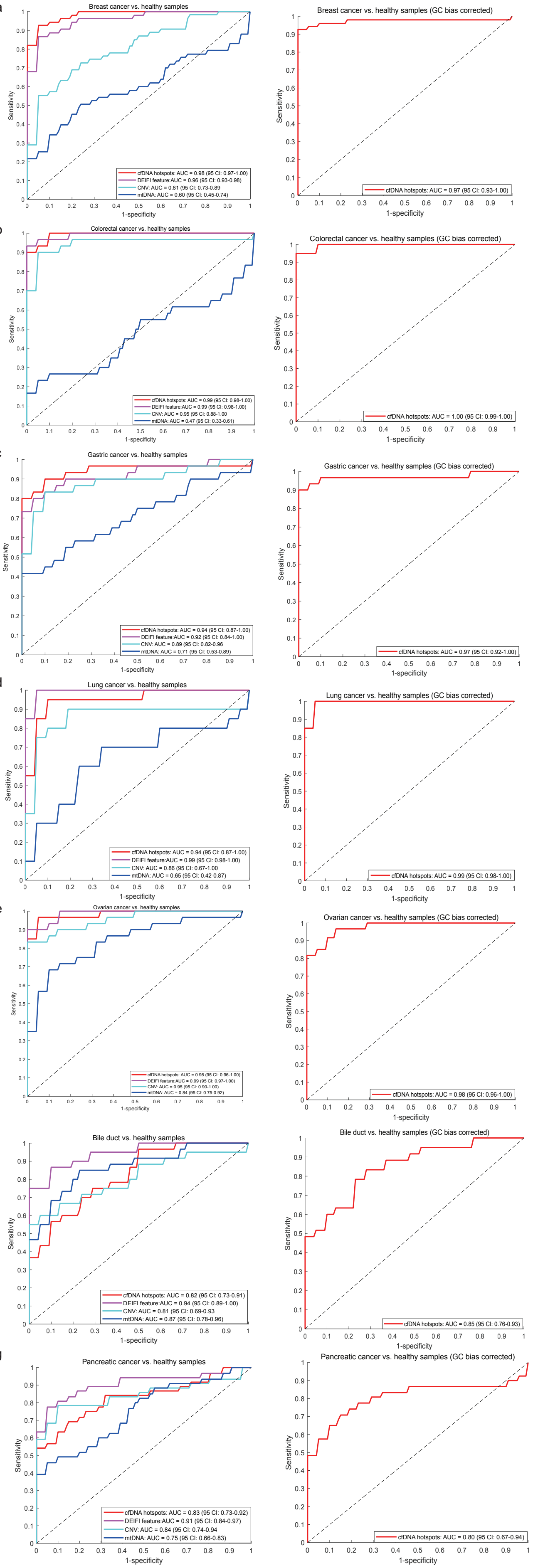

**Fig. S15. Receiver operator characteristics (ROC) for the detection of different early-stage cancers by using IFS from cfDNA fragmentation hotspots before (left panel) and after (right panel) GC bias correction**. (a). Breast cancer. (b). Colorectal cancer. (c). Gastric cancer. (d). Lung cancer. (e). Ovarian cancer. (f). Bile duct cancer. (g). Pancreatic cancer.

#### Supplementary Figure 16

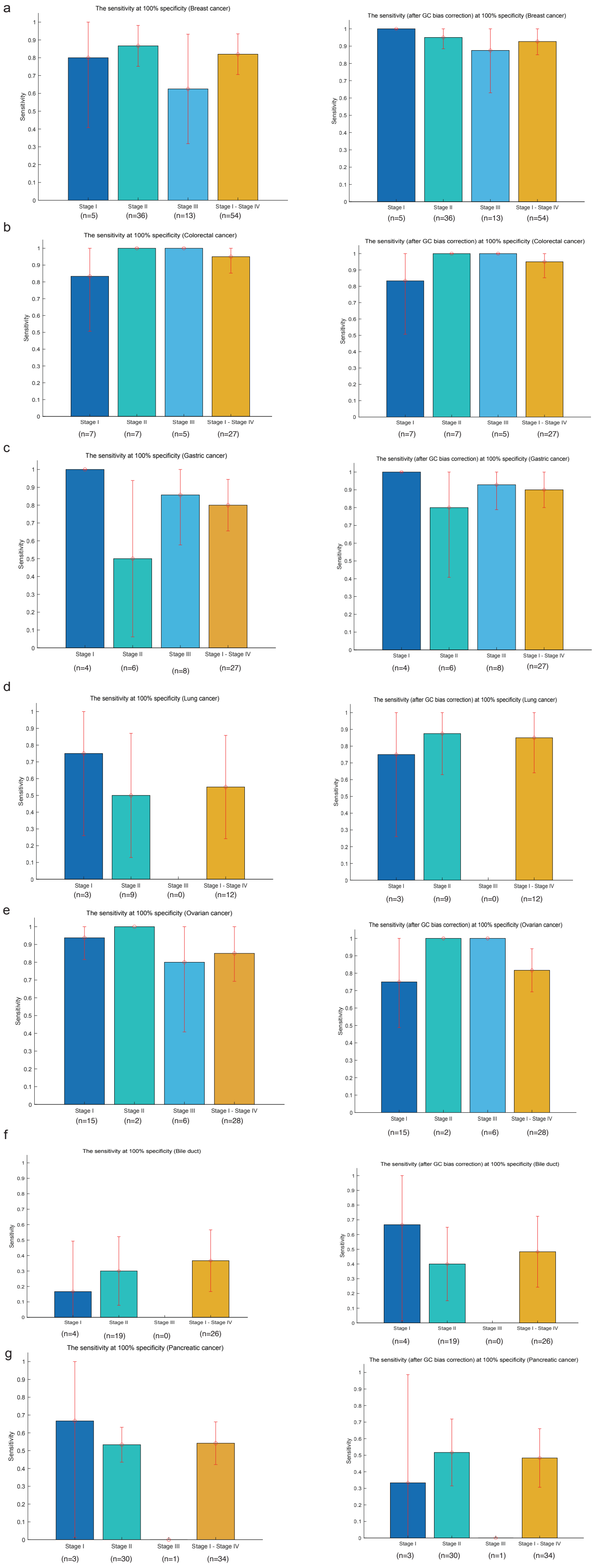

**Fig. S16. The sensitivity across different cancer stages at 100% specificity for the detection of different early-stage cancers by using IFS from cfDNA fragmentation hotspots before (left panel) and after (right panel) GC bias correction**. (a). Breast cancer. (b). Colorectal cancer. (c). Gastric cancer. (d). Lung cancer. (e). Ovarian cancer. (f). Bile duct cancer. (g). Pancreatic cancer. Error bars represent 95% confidence intervals. The sample size in each stage is at the bottom of each bar.

**Supplementary Figure 17**

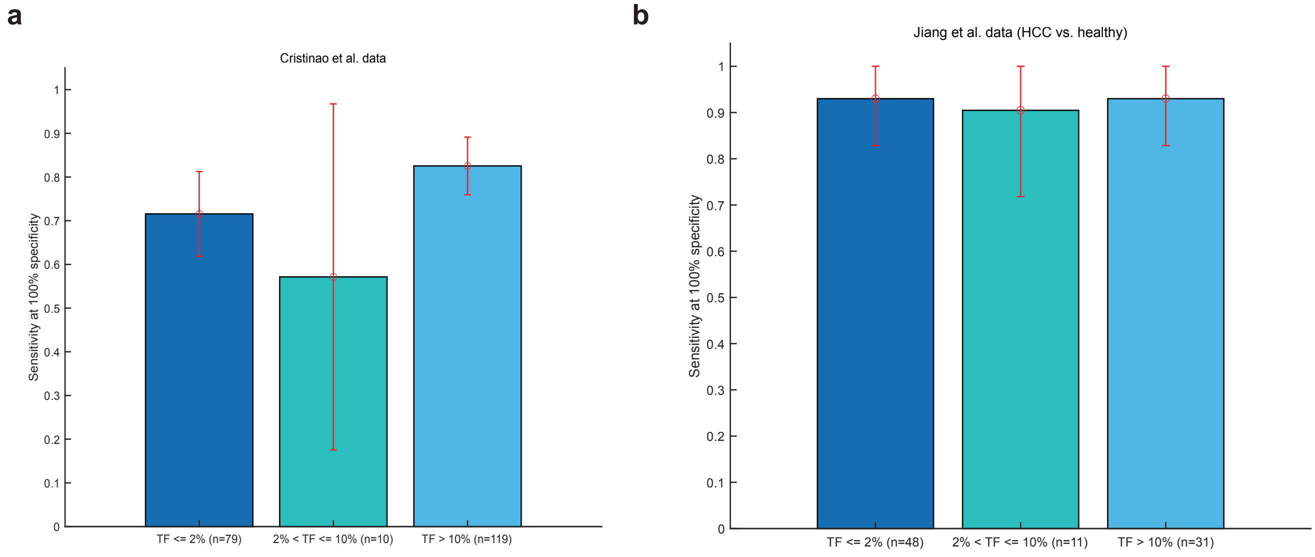


**Fig. S17. The sensitivity at 100% specificity for the detection of early-stage cancer across different tumor fractions**. (a) Cristiano et al. data and (b) HCC vs. Healthy at Jiang et al. data. The tumor fraction is estimated by ichorCNA.

**Supplementary Figure 18**

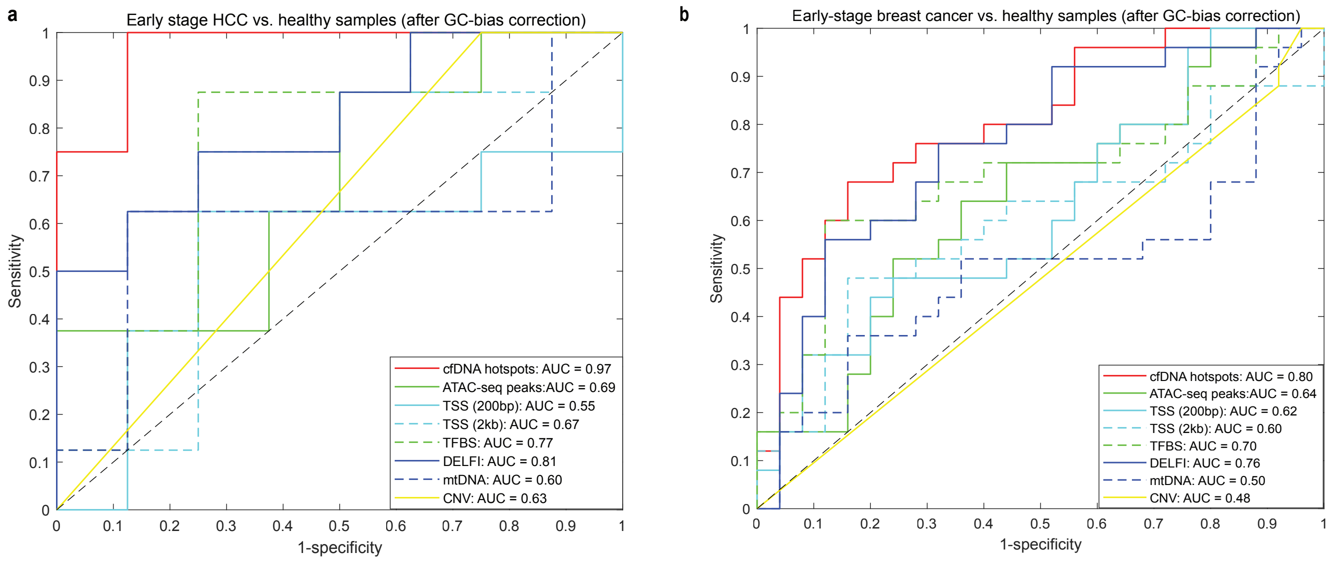


**Fig. S18. Receiver operator characteristics (ROC) at the independent early stage (a). HCC and (b). breast cancer.** Hotspots and machine learning models were identified and trained by Jiang et al. 2015 data (HCC) and Cristiano et al. 2019 data (breast cancer), respectively.

### Supplementary Figure 19

a.

b.

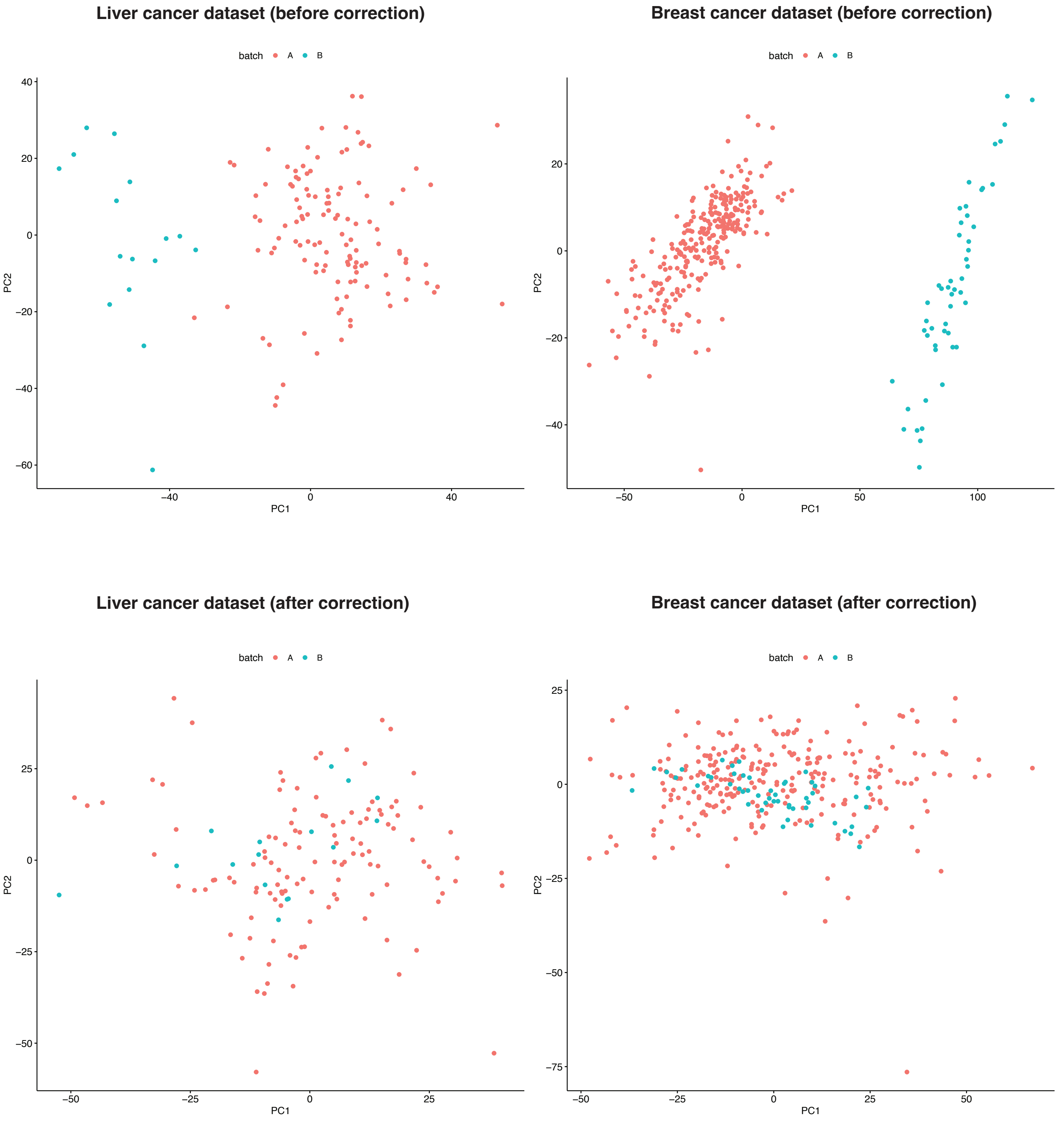


**Fig. S19. PCA results on the IFS Z-score at fragmentation hotspots before (a) and after (b) batch effect correction.** Batch A is the public dataset (Breast cancer and its healthy controls are from Cristiano et al. 2019, Liver cancer and its healthy controls are from Jiang et al. 2015 PNAS). Batch B is our internal test dataset.

### Supplementary Figure 20

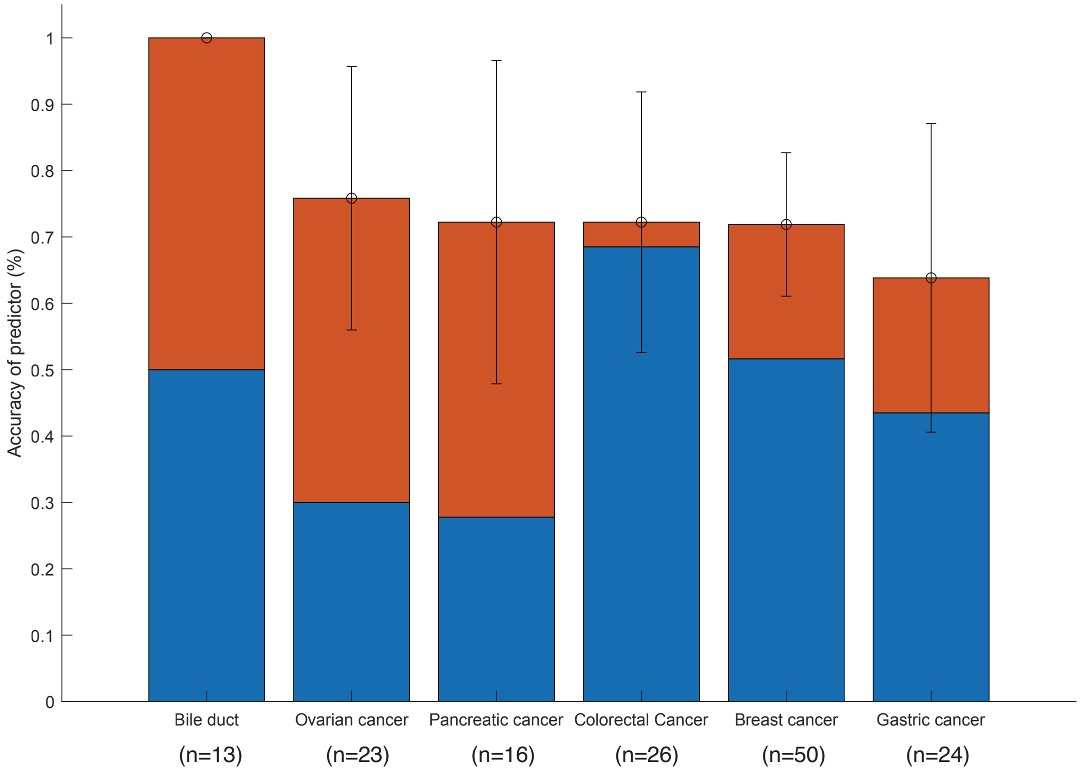


**Fig. S20. Tissues-of-origin prediction across six different cancer types**. Percentages of patients correctly classified by one of the two most likely types (sum of orange and blue bars) or the most likely type (blue bar). Error bars represent 95% confidence intervals.

### Supplementary Figure 21

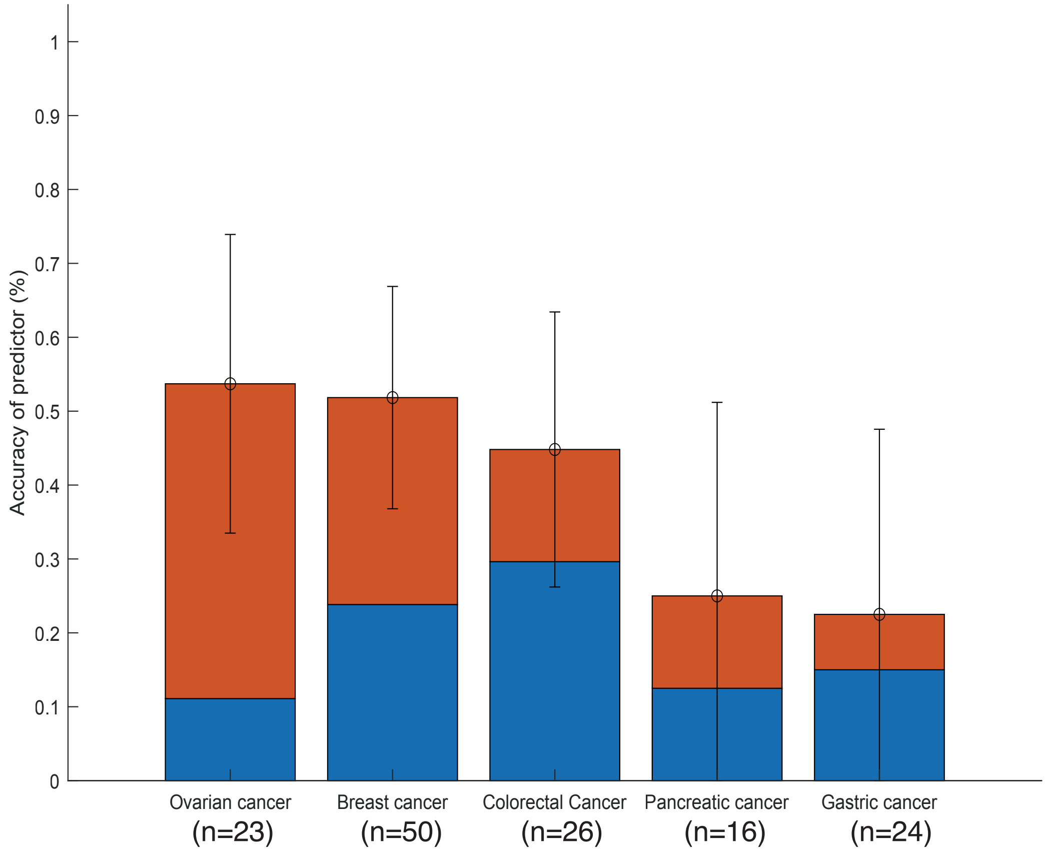


**Fig. S21. Tissues-of-origin prediction randomly by sample frequency across five cancer types**. Percentages of patients correctly classified by one of the two most likely types (sum of orange and blue bars) or the most likely type (blue bar). Error bars represent 95% confidence intervals.

Table. S1. List of public datasets used in the study.

Table. S2. Significantly differentiated fragmentation hotspots between early-stage HCC and Healthy.

Table. S3. Performance evaluation for the detection of early-stage HCC (before GC bias correction).

Table. S4. Performance evaluation for the detection of early-stage HCC (after GC bias correction).

Table. S5. Performance evaluation to distinguish early-stage HCC with benign conditions (HBV-associated liver cirrhosis and chronic HBV infection).

Table. S6. Performance evaluation to distinguish early-stage HCC with benign conditions (HBV-associated liver cirrhosis and chronic HBV infection) (after GC bias correction).

Table. S7. Performance evaluation for the detection of multiple cancer types (after GC bias correction).

Table. S8. Performance evaluation for the detection of multiple cancer types (before GC bias correction).

Table. S9. Patients' information for the independent validation sets.

Table. S10. Performance evaluation at the independent validation sets (after GC bias correction).

Table. S11. Performance evaluation with a fixed cut-off at both training and test set (after batch effect correction).

Table. S12. Performance evaluation for the localization of five or six cancer types.

Data file S1. Zipped source code and readme files. Available in GitHub and Zenodo.org (https://doi.org/10.5281/zenodo.5227520).

Data file S2. The collections of cfDNA fragmentation files and their hotspots identified at different conditions in the study. Available in Zenodo.org (https://doi.org/10.5281/zenodo.5227520).
